# The stress-sensing domain of activated IRE1α forms helical filaments in narrow ER membrane tubes

**DOI:** 10.1101/2021.02.24.432779

**Authors:** Stephen D. Carter, Ngoc-Han Tran, Ann De Mazière, Avi Ashkenazi, Judith Klumperman, Grant J. Jensen, Peter Walter

## Abstract

The signaling network of the unfolded protein response (UPR) adjusts the protein folding capacity of the endoplasmic reticulum (ER) according to need. The most conserved UPR sensor, IRE1α, spans the ER membrane and activates through oligomerization. IRE1α oligomers accumulate in dynamic foci. We determined the *in-situ* structure of IRE1α foci by cryogenic correlated light and electron microscopy (cryo-CLEM), combined with electron cryo-tomography (cryo-ET) and complementary immuno-electron microscopy. IRE1α oligomers localize to a network of narrow anastomosing ER tubes (diameter ~28 nm) with complex branching. The lumen of the tubes contains protein filaments, likely composed of linear arrays of IRE1α lumenal domain dimers, arranged in two intertwined, left-handed helices. Our findings define a previously unrecognized ER subdomain and suggest positive feedback in IRE1 signaling.

## Introduction

The vast majority of secreted and transmembrane proteins mature in the endoplasmic reticulum (ER), which provides a specialized folding compartment with unique biochemical and proteomic characteristics (*1*). To ensure proper folding, a network of quality control pathways, collectively termed the unfolded protein response (UPR), continuously monitors ER folding status and adjusts its capacity according to cellular demand (*2*). The three branches of the metazoan UPR, named after their pivotal ER-resident sensors (IRE1 (*3*, *4*), PERK, and ATF6), activate in response to disruptions in protein folding homeostasis, characterized by an accumulation of unfolded proteins within the ER lumen. This condition, known as ER stress, triggers corrective cellular measures or, if the defect cannot be corrected, apoptotic cell death (*5*).

The life/death decision made after UPR activation involves a molecular timer in which IRE1 activation initially provides cytoprotective outputs but then attenuates even under conditions where ER stress remains unmitigated (*6*). IRE1 attenuation thus predisposes the cell to apoptosis as a consequence of persistent, unopposed PERK signaling (*7, 8*). Therefore, IRE1 signaling in particular and UPR signaling in general sit at a junction of cellular homeostasis and cell death. Maladaptive UPR signaling is a hallmark of many diseases, including cancer, diabetes, and neurodegeneration (*9*).

Mammalian IRE1 has two paralogs (*10*), IRE1α and IRE1β. IRE1α is the major isoform expressed in most cell types. It is an ER-transmembrane protein bearing an ER-stress sensing domain on the ER lumenal side and kinase/ribonuclease (RNase) effector domains on the cytosolic side (*3*, *4*, *11*). IRE1α’s lumenal domain (LD) is bound by the ER-lumenal HSP70-like chaperone BiP, which dissociates upon ER stress (*12*). IRE1α-LD then binds directly to accumulated unfolded proteins, which triggers its oligomerization (*13*, *14*). Oligomerization of the LD in turn drives the juxtaposition of IRE1α’s cytosolic kinase/RNase domains, which activate after *trans*-autophosphorylation (*15*). Lipid stress can likewise induce IRE1 activation through oligomerization, bypassing a need for the LD (*16, 17*). IRE1α’s activated RNase domain initiates the non-conventional splicing of its substrate *XBP1* mRNA (*18*). Spliced *XBP1* mRNA is translated to produce the potent transcription factor XBP1s that upregulates hundreds of genes to restore ER homeostasis. A second consequence of IRE1α RNase activation, termed RIDD (regulated IRE1-dependent mRNA decay), is the selective degradation of a spectrum of ER-bound mRNAs (*19*). RIDD reduces the protein folding burden in the ER and initiates other protective effects (*7, 20*), thus synergizing with *XBP1* mRNA splicing to alleviate ER stress and preserve cell viability.

Upon UPR induction, a fraction of IRE1 molecules cluster into large oligomers that are visible as discrete foci by fluorescence microscopy (*21, 22*). This extensive oligomerization is consistent with the observation that IRE1’s lumenal and cytosolic domains form dimers and higher-order oligomers *in vitro*, also observed in various crystal structures (*23–25*). Both oligomerization of the kinase/RNase domains *in vitro* and formation of foci in cells correlate with high enzymatic activity (*25*).

We and others have shown that IRE1α foci are entities with complex morphology and dynamic behaviors (*26, 27*). Foci comprise two distinct populations of IRE1α (*27*). A small fraction of clustered molecules rapidly exchanges with the pool of dispersed IRE1α in the ER membrane, while the majority are diffusionally constrained to a particular cluster and remain there until that cluster’s eventual dissolution. When foci disappear at late timepoints of stress, their constituent IRE1α molecules are efficiently recycled back into the ER network rather than degraded. The foci are therefore not phase-separated liquid condensates but resemble higher-order arrangements whose assembly and disassembly are likely regulated by distinct mechanisms. However, the molecular principles that explain IRE1α’s complex dynamics during ER stress and the functional consequences of its different assembly states remain a mystery.

## Results

### IRE1α oligomers localize to specialized ER regions with complex topology

Leveraging recent advances in *in situ* structural determination of protein complexes in their native cellular environment (*28, 29*), we applied cryo-CLEM to determine the ultrastructure of IRE1α foci in mammalian cells. To this end, we constructed stable cell lines that express fluorescently tagged human IRE1α under control of an inducible promoter (*14, 27*). We grew cells directly on fibronectin- or collagen-coated gold EM grids to 15% confluency and induced IRE1α expression. We next promoted ER stress with tunicamycin, which blocks N-linked protein glycosylation in the ER lumen, thereby activating IRE1α. We added blue fluorescent nanospheres to the grids as positional markers and plunge-froze the samples in a mixture of liquid ethane/propane at 77 K (*30*).

To localize IRE1α foci, we first imaged the frozen grids on a fluorescent light microscope fitted with a liquid nitrogen sample chamber (*31*). In our initial studies with an IRE1α-GFP fusion protein, the stressed cells exhibited strong auto-fluorescence at liquid nitrogen temperature (77 K) as previously observed (*32*), which hindered identification of IRE1α foci (Fig. S1). To overcome this hurdle, we fused IRE1α to the exceptionally bright fluorescent protein mNeonGreen (mNG). This experimental refinement revealed spots emitting high fluorescence in the green but much lower fluorescence in the red and blue channels (Fig.1A). Spots meeting these criteria were absent in control cells expressing IRE1α not fused to mNG (Fig. S1), while auto-fluorescent spots with high green, blue and red signals were abundant in both samples. Thus, plotting the ratio of green/red and green/blue fluorescence intensity allowed us to exclude non-specific signals (Fig. S1) in order to identify IRE1α-mNG foci reliably.

**Figure 1.**
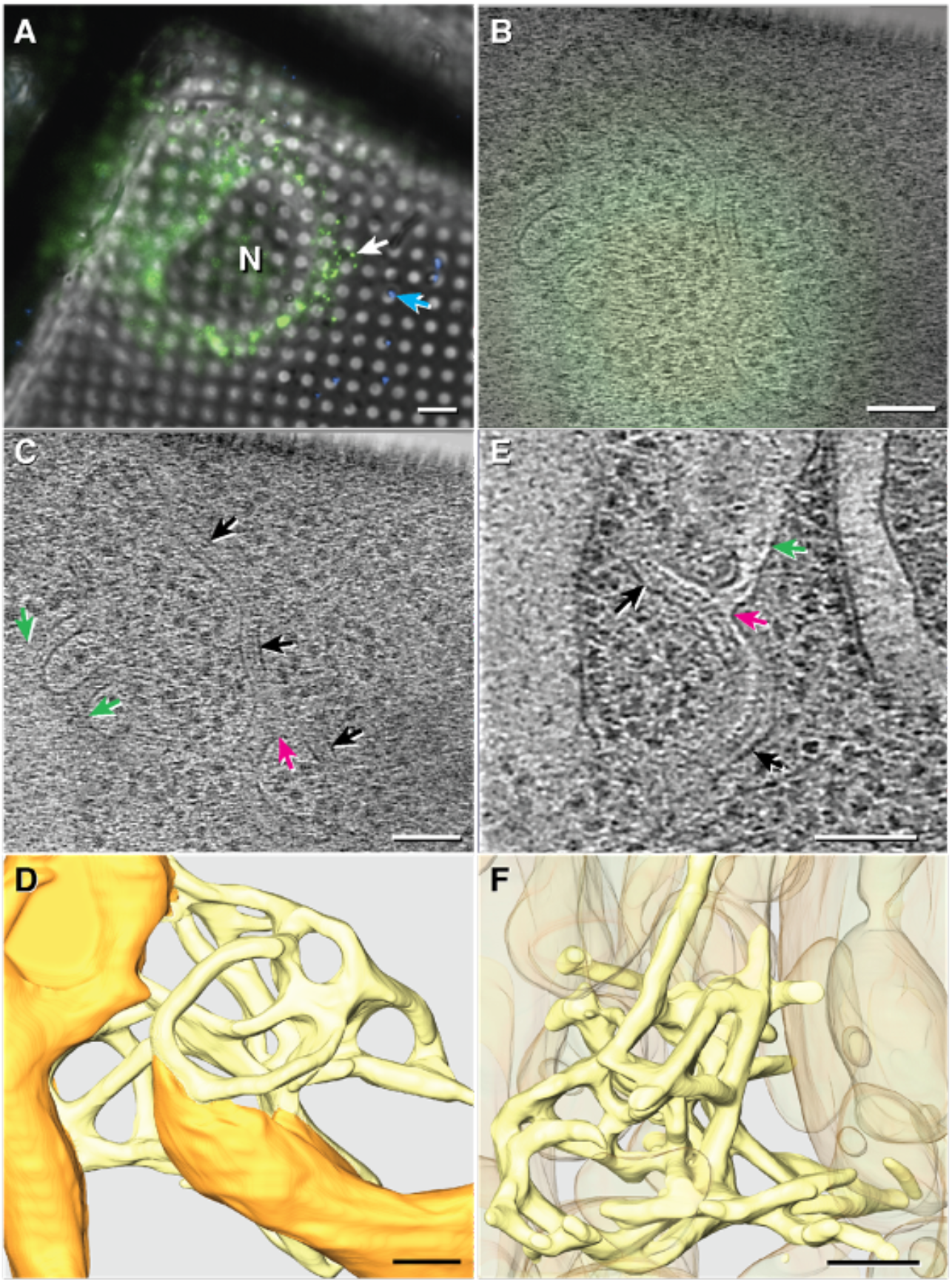
IRE1α oligomers localize to specialized ER regions with complex topology. (A) Fluorescent profiles imaged at liquid nitrogen temperature for stressed cells grown on EM grids and expressing IRE1α-mNG. White arrow denotes the fluorescent spot corresponding to the tomogram depicted in (B-D). Blue arrow: 500 nm fluorescent nanosphere. “N”: Nucleus. Scale bar = 6 μm. (B) Correlation of fluorescence image with a representative z slice (C) of the tomogram showing examples of narrow membrane tubes (black arrows) connected to general ER network at junctions (green arrows) and to each other at three-way junctions (magenta arrows). The tomogram was manually segmented for 3D visualization in (D), where normal ER membranes are depicted in orange and constricted membranes colocalizing with IRE1α-mNG signal are yellow. (E) A representative z slice from a higher resolution tomogram obtained in stressed U2OS-IRE1α-mNG cells. (arrows color code same as (C)). (F) manual segmentation of region shown in (E) with constricted membranes shown in yellow and other membranes shown in orange at 50% transparency. Scale bars for B-F = 100 nm. Densities corresponding to ribosomes and cytoplasmic densities in (D) and (F) are omitted for clarity. See Fig. S4.

We next imaged IRE1α foci by cryo-CLEM combined with cryo-ET. To this end, using nanospheres, grid features, cell boundaries and other landmarks (such as ice), we located the same IRE1α foci with the electron microscope that we had previously identified with the light microscope and then recorded tilt-series. Across 9 tomograms obtained from mouse embryonic fibroblasts (MEFs), fluorescent IRE1α foci consistently localized to specialized regions of the ER that display a network of remarkably narrow, anastomosing tubes (Fig. 1B-D and Fig. S2–3) with an average diameter of 28 ± 3 nm (± standard deviation). As visualized in segmented three-dimensional (3D) reconstructions, the tubes frequently connect with each other by three-way junctions and to surrounding ER structures, forming a topologically complex yet continuous membrane surface (Fig. 1D, Supp Fig. S2–3). Unlike the surrounding ER, the narrow anastomosing tubes colocalized with IRE1α foci are devoid of bound ribosomes (Fig. S4).

To confirm these results and obtain higher resolution images, we used human osteosarcoma U-2 OS cells that likewise express inducible IRE1α-mNG (27). Compared to MEFs, U-2 OS cells spread more and therefore contain expansive thin regions that enhance the resolution of cryo-ET imaging. We imaged tilt series from 8 IRE1α-mNG foci at slightly higher magnification (33,000x for U-2 OS cells vs. 22,000x for MEFs). We again observed thin anastomosing tubes with similar characteristics as those in MEFs, including the absence of bound ribosomes, three-way junctions and connections to adjacent ER (Fig. 1E-F and Fig. S4–5). Using neural network-enhanced machine learning to segment the tomograms (33), we confirmed the basic features of the manual 3D reconstructions without subjective bias (Fig. S6). Taken together, IRE1α foci localize to a highly specialized ER region, henceforth termed the “IRE1α subdomain”.

### Orthogonal methods reveal IRE1α subdomains

To validate our discovery of IRE1α subdomain tubes with alternative approaches, we performed conventional and immuno-electron microscopy on HEK293 cells expressing IRE1α-GFP fusion protein (22). In ER-stressed HEK293 cells, electron micrographs of chemically fixed, stained and Epon-embedded thin sections exhibit thin membrane tubes and networks of comparable topology to those seen in the cryo-tomograms (Fig. S7) and unlike anything seen in un-stressed, control cells. These structures are infrequently observed and stained more strongly in their lumenal space than the surrounding ER, suggesting the presence of a high protein density inside and likely represent IRE1α subdomains.

To directly identify IRE1α foci in electron micrographs, we next performed immunogold labeling of ultrathin cryosections of HEK293 cells expressing IRE1α-GFP. As above, we induced ER stress with tunicamycin to drive activated IRE1α into foci. In non-stressed cells, gold particles specific to IRE1α-GFP sparsely label large regions of the cell with visible ER (Fig. 2A-A’ and Fig. S8). By contrast, in stressed cells we observed clusters of gold particles in regions of much higher membrane complexity (Fig. 2B-B’, 2E-F’ and Fig. S8). In these regions, we observed narrow membrane tubes of similar diameter (~28 nm) as both longitudinal and transverse cross sections. Quantification of the inter-particle distances between samples revealed a clear difference in gold particle density (Fig. 2C-D), reflecting a population of clustered IRE1α - GFP molecules in stressed cells that is absent in non-stressed control cells. This observation is consistent with previous data that show that IRE1α molecules, which uniformly distributed in the ER during homeostasis, aggregate in foci of dozens of IRE1α molecules during ER stress induction (22, 26, 27). Notably, we observed a distinct lumenal density inside the membrane tubes, which is circular in transverse cross sections (Fig. 2E-F’).

**Figure 2.**
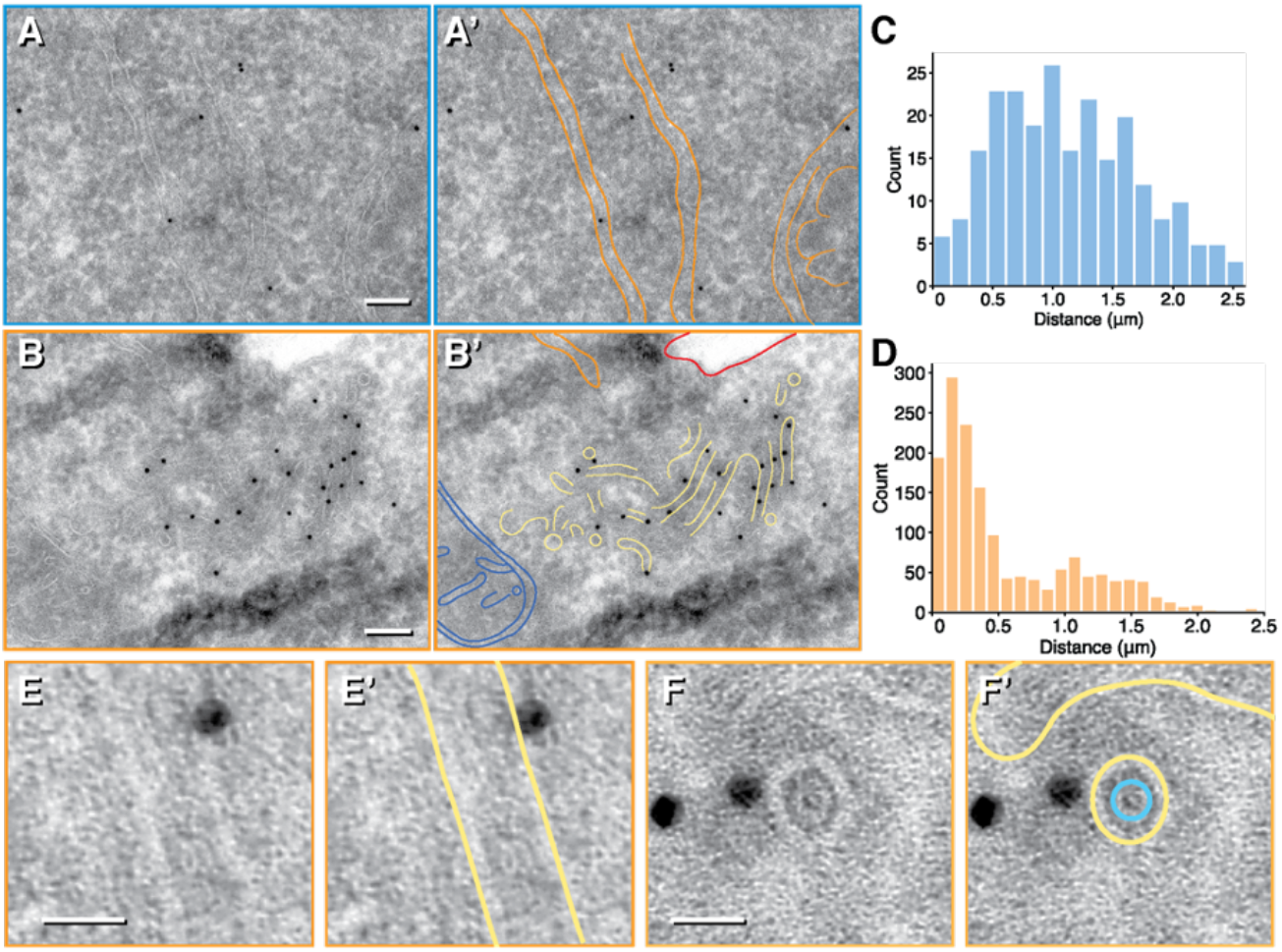
Orthogonal immuno-electron microscopy reveals IRE1α subdomains. (A, A’) Representative micrograph of cells expressing IRE1α-GFP but not subjected to ER stress induction by Tunicamycin (Tm). Gold particles recognizing IRE1α-GFP epitope (via binding to anti-GFP primary antibody) sparsely label general ER structures. Scale bar = 100 nm. In A’, orange: ER sheet/tubule membranes (B, B’) Representative micrograph of cells treated with Tm where gold particles recognizing IRE1α-GFP epitope densely localizes to a region enriched in narrow membranes of 26 ± 2 nm diameter. Scale bar = 100 nm. In B’, blue: mitochondrion, orange: ER sheet/tubule membranes, red: plasma membrane, yellow: narrow IRE1α subdomain membranes. Histograms of inter-gold particle distances measured in micrographs from non-stressed samples (C) and stressed samples (D) reveal a population of densely clustered gold particles enriched with ER stress induction. Zoomed in view showing longitudinal (E, E’) and end-on (F, F’) tube cross-sections with ~28 nm diameter close to gold particles. Scale bar = 20 nm. A ring-like density within the lumenal space is clearly visible in F (segmented in teal)

### IRE1α subdomain membrane tubes contain lumenal helical filaments

Consistent with the density observed in immunogold labeling experiments, we observed regular densities in the lumen of the IRE1α subdomain tubes in cryogenic tomographic reconstructions, which we interpret as oligomers of IRE1α-LD. In our MEF-derived tomograms, the lumenal densities resemble train tracks that in longitudinal sections run parallel to the membranes (Fig. 3A-A’). Closed rings roughly concentric with the enclosing tube membrane are clearly visible in instances where IRE1α subdomain tubes are imaged parallel to the beam direction (Fig. 3B-B’ and Fig. S9). The inner rings measure 9 ± 0.5 nm in diameter and are enclosed by membrane tubes that are approximately 28 ± 1 nm diameter (Fig. 3C).

**Figure 3.**
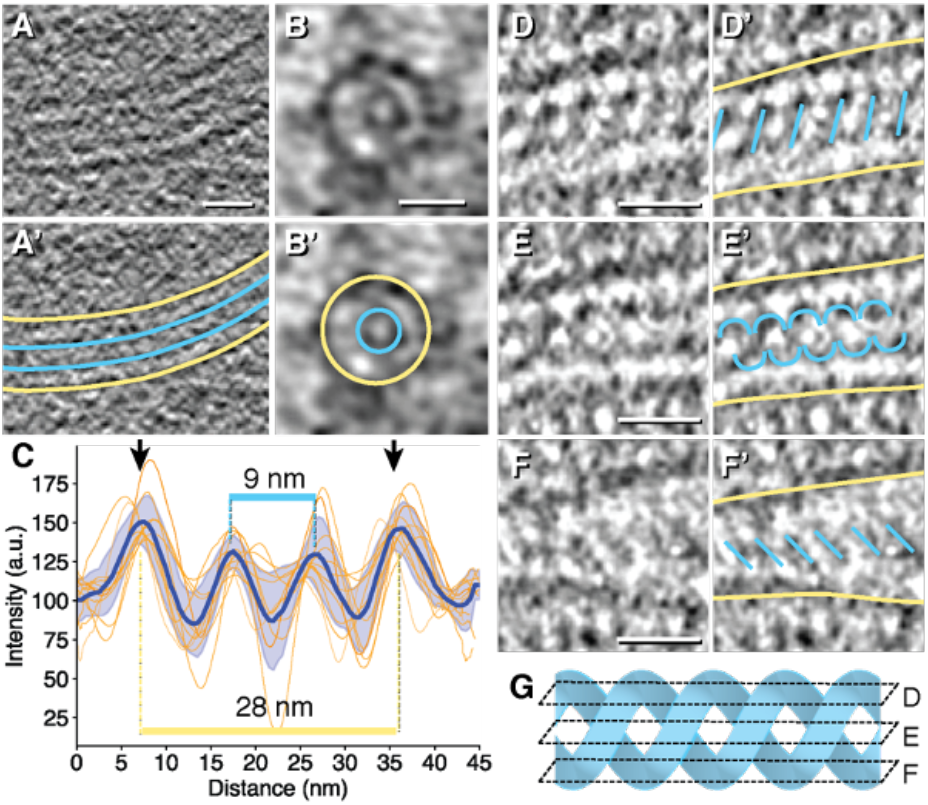
IRE1α subdomain membrane tubes contain lumenal protein densities. Representative examples of a longitudinal cross-section (A, A’) and an end-on cross-section (B, B’) obtained in MEFs-IRE1-mNG cells revealing membrane density (yellow) surrounding lumenal protein density (teal). Scale bar = 20 nm. Intensity line plots across subdomain tubes are aligned, plotted and averaged across 9 cross-sections and plotted as a function of distance in (C). Blue line with shaded error of the mean is the averaged trace for all plots. Distance separation of peak maxima are indicated for peaks representing membrane densities (yellow) and those representing protein densities (teal). (D-F’) show an example of lumenal protein densities with helical features obtained in U2OS-IRE1-mNG cells viewed as top (D, D’), middle (E, E’) and bottom (F, F’) sections. Scale bars = 20 nm. G) Schematic of an idealized double helix to illustrate how the cryo-tomogram slices in D-F would intercept as planes and give rise to the corresponding densities segmented in D’-F’.

Strikingly, in the tomograms of U-2 OS cells, the lumenal densities show sufficient substructure to reveal two intertwined helices (Fig. 3D-F’). This helical feature is most clearly seen in IRE1α subdomain tubes in which the top and bottom cross sections show equidistant parallel angled lines of opposite directionalities, whereas the middle cross section shows helical features (Fig. 3D-G).

### Sub-tomogram averaging resolves flexible IRE1α-LD double helices

To determine the 3D structure of the lumenal double-helical density, we extracted subvolumes along the membrane tubes for subtomogram averaging, using the EMAN2 tomogram processing workflow (Fig. S10). The resulting electron density map obtained by averaging of 653 subvolumes portrayed a double helix composed of two equidistant intertwined strands with a pitch of 17 nm in each individual strand (Fig. 4A, B and Fig. S10). Analysis of sequential cross sections revealed that the double helices are left-handed (Fig. 4A). The handedness was confirmed by subtomogram averaging of ribosomes in the same tomograms (data not shown). This double helical structure is reminiscent of the unit cell of the *S. cerevisiae* IRE1-LD crystal structure ((*23*) PDB ID: 2BE1), in which two-fold symmetrical head-to-head IRE1-LD dimers arrange into helical filaments by forming tail-to-tail contacts. Two such yeast IRE1-LD filaments intertwine in left-handed double helices.

**Figure 4.**
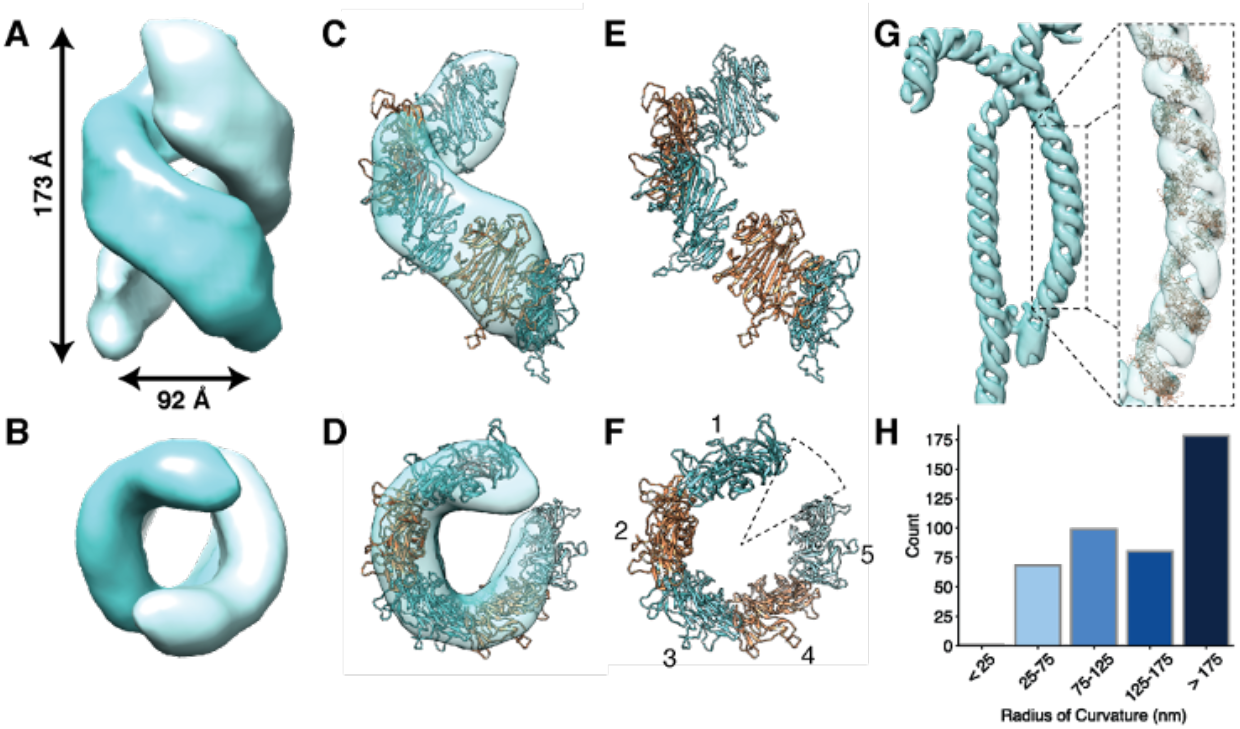
Sub-tomogram averaging resolves flexible IRE1α-LD double helices. (A-B) Electron density map obtained by sub-tomogram averaging of 653 subvolumes in U2OS-IRE1-mNG tomograms with indicated distances for helical pitch and width. (C, D) Semi-transparent masked average of one strand of the double helix fitted with modeled IRE1α lumenal oligomer (E, F) using Chimera’s “fit-in-map” function. 77% of the structure fit within this density. (B, D and F) are rotation of (A, C, E) by 90 degree along × axis. (F) 5 dimers of IRE1α-cLD are fitted into masked single-strand map, but approximately 6 dimers are required to complete one full turn. (G) An isosurface of the averaged density mapped back on to the cryo-tomogram at a highly curved region and fitted with a helix of the modeled IRE1α lumenal structure using a range of interface angles to accommodate curvature. (H) A histogram of measured radii of curvature observed for 25 nm segments along IRE1α subdomain network.

With fluorescence, immunogold labeling, and structural (left-handed intertwined helices) evidence that the regular density in the tubes is IRE1α-LD, we endeavored to interpret its structure in light of existing atomic models. The two most relevant structures are (i) the crystal structure of the active, polymeric yeast IRE1-LD, which forms intertwined left-handed helices like those in the cryo-tomograms but with different pitch (38 nm); and (ii) the crystal structure of the human IRE1α-LD in its inactive dimeric form. We chose to begin with the former and used SWISS-MODEL homology modeling to predict a homologous human IRE1α-LD structure based on the yeast structure (Fig. S11) (*34*). The resulting model retains similarity in secondary and tertiary structure compared to the (inactive) human IRE1α-LD but contains rearranged elements at the tail-ends of the homodimer (Fig. S11–12). We next fit the modeled head-to-head IRE1α-LD dimer into our double-helical map as a rigid body and modified the tail-to-tail dimer interface so that the polymer would exhibit the same helical pitch observed in the tomograms (Fig. 4C-D and Figs. S13). We maintained both head-to-head and modified tail-to-tail interfaces in propagating the array of dimers to occupy the helical map. The resulting model contains approximately six dimers per turn in each helical filament (Fig. 4E-F), consistent with the *S. cerevisiae* LD crystal structure (Fig. S14) but with a compressed pitch (17 nm vs. 38 nm in the yeast crystal structure). We conclude that an IRE1α-LD polymer accounts well for the regular density, including in the details of its thin ribbon-like shape. By analogy to the yeast LD structure, we fit the human LD into the density with its putative unfolded protein binding groove facing towards the membrane; however, at the current resolution this assignment must be considered tentative because no secondary structural features are resolved that would support this orientation. Likewise, the details of the subunit contacts remain uncertain.

Mapping the averaged subvolumes back onto the cryo-tomograms (Fig. 4G) allowed us to generate a volumetric map for IRE1α-LD within an IRE1α subdomain. The resulting distribution of double-helical filaments showed that the helices are not stiff but curve to varying degree (Fig. 4H and Fig. S15), ranging from straight segments (radius of curvature > 175 nm) to segments bent with radii approaching 25 nm. This bending indicates that IRE1α-LDs, their oligomerization interfaces, and/or inter-strand connections must undergo conformational rearrangements that can accommodate the observed range of curvatures.

### IRE1α-LD helices and IRE1α subdomains are flexible

Further examination of the tomograms revealed infrequent instances in which lumenal double helices are irregularly spaced from the tubular membranes (Fig. S16). In one case, we observed helices that are not completely enclosed by membrane tubes but instead are positioned on the lumenal face of a flat ER membrane (Fig. 5A-B’).

**Figure 5.**
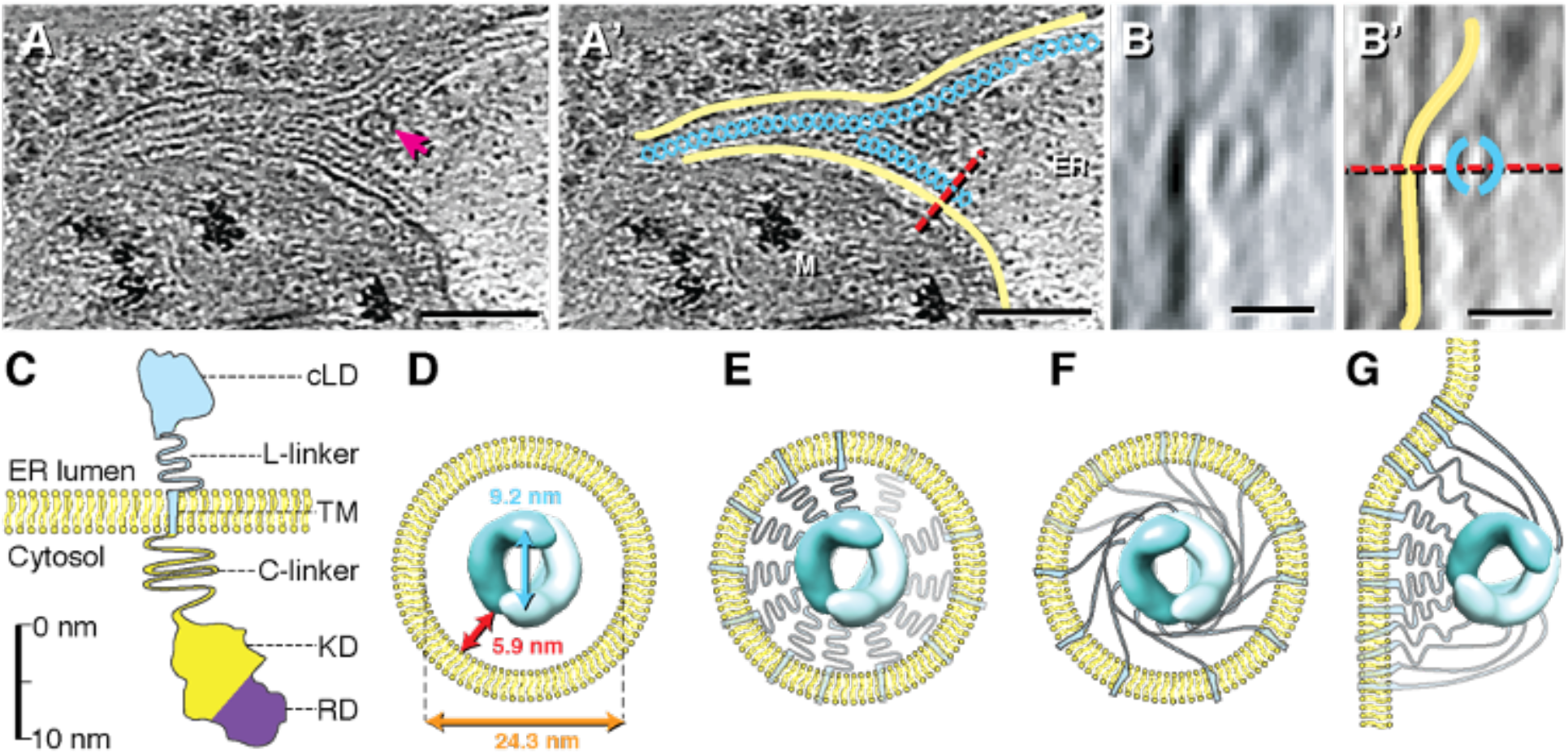
IRE1α-LD helices can accommodate a range of distance from membrane. (A, A’) An instance of ordered IRE1α-LD helices not completely enveloped by narrow membrane tubes. Arrow in A indicates a membrane fenestration. ER: lumenal space; M: mitochondrion. Scale bar = 100 nm. (B, B’) Side view along indicated plane obtained by 90-degree rotation along × axis. Scale bar = 20 nm. (C) Diagram of IRE1α domain architecture drawn to approximate scale. cLD: core lumenal domain (a.a. 19-390), L-linker: lumenal linker (a.a. 391-443), TM: transmembrane helix (a.a. 444-464), C-linker: cytoplasmic linker (a.a. 465-570), KD: kinase domain (a.a. 571-832), RD: ribonuclease domain (a.a. 835-964). Scale bar = 10 nm. (D) Dimensions of IRE1α-LD helices within IRE1 subdomain lumenal space. (E, F) Schematics for alternative TM domain and L-linker arrangements within the narrow subdomain tubes. There are 24 monomers per turn for the double helices, but only 12 TM and L-linker domains are shown for clarity. (G) Model for TM and L-linker arrangement for helices not completely surrounded by membrane as seen in B

The helically arranged IRE1α core-LDs (cLD) are each attached via a 52-amino acid linker to IRE1α’s transmembrane domains, which in turn are connected to the cytosolic kinase/RNase domains (Fig. 5C). In IRE1α subdomain tubes, the linker domain could either be in a compacted or an extended conformation to bridge the 5.9 nm distance between the helix and the membrane (Fig. 5D; and Fig. 5E-F, respectively). By contrast, when IRE1α-cLD helices are attached to a flat ER membrane surface on one side only as shown in Fig. 5A-B, their linker domains must accommodate the different distance requirements imposed by the positioning of individual IRE1α-cLD monomers to reach the nearest membrane surface, likely conforming to a range of compaction (Fig. 5G). One consequence of this arrangement is that IRE1α kinase/RNase domains on the cytosolic side of the membrane are brought into even closer proximity and hence experience a higher local concentration (we estimate it likely exceeding 1 mM, see Methods) than in the tubes, in which the membrane surface surrounding the helices is ~4-fold larger than the contact surface of the helices attached to a flat membrane sheet.

## Discussion

The ER is formed from a single continuous membrane that is dynamically differentiated into a plethora of pleomorphic subdomains (*35*), including the nuclear envelope, smooth tubules, tubular matrices (*36*), ribosome studded flat sheets, ER exit domains, inter-organellar contact sites (*37*), and ER-phagic whorls (*38*). Using cryo-CLEM to inspect IRE1α fluorescent puncta at macromolecular resolution, we found that UPR activation leads to the formation of previously unrecognized “IRE1α subdomains”, comprised of topologically heterogeneous assemblies of anastomosing ~28 nm membrane tubes. Inside the highly constricted lumenal spaces of this labyrinthine network, IRE1α-LDs assemble into ordered double-helical filaments. Our use of cryo-CLEM and cryo-ET to visualize IRE1α oligomers directly in intact cells demonstrates the power of *in situ* structural biology, providing insight into the supramolecular arrangements of molecules in their native environment.

Four independent lines of evidence support our conclusion that the observed helical densities indeed correspond to oligomerized IRE1α-LDs: (i) In 20 out of 20 fluorescent foci analyzed by EM, we observed narrow membrane tubes, which in 18 out of 20 have a consistent diameter of ~28 nm and contain lumenal protein density consistent with the helical reconstruction. By contrast, no such structures were observed in adjacent and random regions of the cell, including those emitting high autofluorescence; (ii) orthogonal analysis by immunogold-staining of thin sections revealed IRE1α localization to similarly-narrow tubes; (iii) the double-helical architecture closely resembles that observed in the crystal structure of yeast IRE1-cLD (*23*); and (iv) the reconstructed helical volume has the same flat ribbon-like shape and dimensions as IRE1α-cLD oligomers.

The accumulation of IRE1α within these specialized structures explains recent observations that IRE1α foci contain a readily exchanging periphery and a diffusionally constrained core (*27*). We surmise that the two distinct populations represent (i) IRE1α molecules located at the subdomain junctions where the 28 nm tubes merge with the main ER and (ii) those located deeper in the interior of the narrow tubes. IRE1α molecules at helix ends located near the tube mouths can readily dissociate, and new IRE1α molecules can associate; they thus represent a freely exchanging pool at the foci’s periphery. By contrast, IRE1α molecules at the foci’s core are physically trapped in regularly arrayed helices; they thus represent a non-exchangeable pool.

The confinement of IRE1α in the anastomosing IRE1α subdomains suggests functional consequences for the regulation of IRE1 signaling. The presence of just a single unfolded protein molecule in each 100 nm-long cylinder segment amounts to an effective concentration of ~40 μM, which is in the same order of magnitude as the affinity measured for IRE1-unfolded protein binding *in vitro* (*13, 14*). Thus a few unfolded protein molecules trapped inside the tubes would saturate IRE1α LD with activating ligand, triggering a positive feedback loop that effectively locks IRE1α into its activated state. This effect is due to the enormous concentration IRE1α experiences upon foci formation. Without UPR activation, IRE1α-mNG is distributed over the ER surface at about 10 molecules per μm^2^ (*27*). By contrast, inside IRE1α subdomains, it is enriched 1500-fold to 15,000 molecules per μm^2^ of ER membrane (Methods). Moreover, based on decreased diffusional freedom due to the complex membrane architecture (*39*) and IRE1α LD’s helical assembly, IRE1α subdomains stabilize the oligomeric state. In an IRE1α subdomain, we estimate that the local concentration of IRE1α’s LD inside the tubes and IRE1α cytosolic domains on the tubes’ cytosolic surface approaches 5 mM and 200 μM, respectively. These concentrations well-exceed the range of affinities measured *in vitro* for purified IRE1 domain oligomerization (*25, 40*).

The resemblance to the crystal structure of the yeast IRE1 cLD is remarkable: this structure likewise reveals two intertwined, equidistant left-handed helices with 12 LDs per turn and strand, albeit 2.2-fold more stretched out along the central axis. It will certainly be interesting to repeat this work with yeast cells to see if the differences are species-specific or due to artefacts of crystallization conditions. Given that two helices of the IRE1α-LDs are equidistantly intertwined, there may be bridges between them, perhaps formed from unresolved regions of the cLD or linker domains, and/or bound unfolded protein chains. We find it equally remarkable that IRE1α helices are observed both in the narrow membrane tubes of the IRE1α subdomain as well as, albeit more rarely, lying flat on the lumenal side of ER sheets. Thus, the lumenal IRE1α linker domains must accommodate a range of distances separating IRE1α-cLD from the membrane surface. These two topologically distinct arrangements of IRE1α-cLD helices may co-exist or could be temporal precursors of one another.

IRE1α subdomains could form by IRE1α-LD filaments pushing finger-like ER membrane protrusions, which then fuse with adjacent ER, or from IRE1α-LD filaments pushing ridge-like deformations into flat ER membranes, which then separate from the flat ER following membrane fission/fenestration (*41*). Alternatively, IRE1α filament polymerization could constrict existing ER tubes to form IRE1α subdomains or stabilize pre-existing narrow membrane tubes (*42*) into subdomain tubes with regular ~28 nm diameter. In support of the latter model, in 2 out of 20 of our tomograms, we observed strong IRE1α-mNG signal in regions with thin and irregular membrane tubes, but no ordered lumenal filaments (Fig. S16). Such structures may represent an intermediate state, captured after IRE1α’s preferential localization but preceding helix formation.

Our discovery of the IRE1α subdomain raises intriguing possibilities with regard to how recruitment of IRE1α into these highly specialized ER structures could serve regulatory functions in the UPR. Its high concentration into long-lived topologically distinct structures may scaffold assembly of downstream effectors, as previously proposed (*9*), and/or affect the selection of IRE1α mRNA substrates in switching between *XBP1* splicing and RIDD activities (*19*). Such regulation may thus profoundly affect the life/death decision that the UPR makes in response to a breakdown in ER protein folding homeostasis and hence be of crucial importance in designing UPR-centered therapies in protein folding disorders.

## Acknowledgements

We thank Margaret Elvekrog, Ariane Briegel, Diego Acosta-Alvear, Richard Fetter, Suzanne van Dijk, Stefan Niekamp and Vladislav Belyy for their advice and technical assistance. We thank Greg Huber, Robert Ernst, Adam Frost and Jodi Nunnari for insightful discussions.

## Author contributions

AA, JK, PW and GJ, conceived of the study. NHT, SDC, PW and GJ designed cryo-CLEM experiments. NHT and SDC generated and screened grids. SDC performed the cryo-CLEM/cryo-ET and subtomogram averaging and generated IRE1α subdomain models, neural network-derived segmentations and movies. NHT performed manual segmentation, quantification of IRE1α subdomain dimension and calculations. ADM performed Epon-embedded electron microscopy and immunogold labeling experiments. NHT and SDC prepared figures. NHT, SDC, PW and GJ wrote the manuscript.

## Data and materials availability

The codes used for data quantification and plots in this paper are freely available at https://github.com/han-tran/IRE1. The final sub-tomogram averaged maps and representative tomograms can be accessed at EMDB entry ID EMD-23058. All raw and processed data are available upon request. All cell lines and plasmids used in this paper are in the Walter lab depository and are available upon request.

## Materials and Methods

### Cell culture and grid preparation

Previously described MEFs-IRE1-mNG and U2OS-IRE1-mNG cell lines (*14, 27*) were cultured in high glucose DMEM media supplemented with 10% tetracycline-free fetal bovine serum (FBS; Takara Bio), 6 mM L-glutamine, and 100 U/ml penicillin/streptomycin. All cell lines used in the study tested negative for mycoplasma contamination when assayed with the Universal Mycoplasma Detection Kit (ATCC 30-1012K). To minimize autofluorescence, the same culture media without phenol red was used to grow cells for grid preparation (*32*). Prior to cell plating, gold Quantifoil London finder grids (EMS R2/2 LF-Au-NH2) were UV treated to sterilize and coated with cell adhesion matrix. For MEFs-IRE1-mNG cell line, grids were coated in droplets of 500 μg/mL Fibronectins (Sigma-Aldrich S5171-.5MG) for 5 minutes on each side, washed in PBS, blotted and air-dried. For U2OS-IRE1-mNG cell line, grids were coated with ~4 mg/mL undiluted Collagen type I (Corning 354236) droplets for 20 minutes, washed in PBS, blotted and air-dried. Cells were seeded at 15% confluence and allowed to adhere for 8 hours and induced with Doxycycline (500 nM) for 6 or 18 hours for MEFs and U-2 OS cell lines, respectively. ER stress was then induced by treatment with 1.5 μg/mL of Tunicamycin for 2 hours.

Immediately prior to being plunge frozen, 3 μl of a beads suspension was added to the grids. The bead suspension was made by a 1:1 dilution of 500 nm blue (345/435 nm) polystyrene fluorospheres (Phosphorex) with a 3:1 concentrated solution of 20 nm:5 nm colloidal gold (Sigma Aldrich) blocked with bovine serum albumin. The gold beads served as fiducial markers for cryo-tomogram reconstruction while the blue fluorospheres served as landmarks for registering cryogenic fluorescence microscopy images collected from different channels as well as with cryo-EM projection images for cryo-CLEM (32). Residual media and bead suspension were blotted manually from the back side with Whatmann paper #40 in 90% humidity. Grids are plunge-frozen in liquid ethane/propane mixture using a Vitrobot Mark IV (FEI, Hillsboro, OR). Plunge-frozen grids were subsequently loaded into Polara EM cartridges (FEI) (*31*). Cryo-EM cartridges containing frozen grids were stored in liquid nitrogen and maintained at ≤−150 °C throughout the experiment including cryogenic fluorescence microscopy imaging, cryo-EM imaging, storage and transfer.

### Fluorescence imaging and image processing

The EM cartridges were transferred into a cryo-FLM stage (FEI Cryostage), modified to hold Polara EM cartridges (*31*), and mounted on a Nikon Ti inverted microscope. The grids were imaged using a 60X extra-long-working-distance air-objective (Nikon CFI S Plan Fluor ELWD 60X NA 0.7 WD 2.62–1.8 mm).

Images were recorded using a Neo 5.5 sCMOS camera (Andor Technology, South Windsor, CT) using a 2D real-time deblur deconvolution module in the NIS Elements software from AutoQuant (Nikon Instruments Inc., Melville, NY). The 2D real-time deconvolution algorithm estimates a PSF using several factors such as sample thickness, noise levels in the image, background subtraction and contrast enhancement. All fluorescence images (individual channels) were saved in 16-bit grayscale format. IRE1α-mNG was visualized with a FITC filter. Blue fluorospheres were visualized with a DAPI filter. Red autofluorescence was imaged using an mCherry filter.

### EM imaging

Cryo-EM grids previously imaged by cryo-LM were subsequently imaged by electron cryo-tomography using a FEI G2 Polara 300 kV FEG TEM (FEI) equipped with an energy filter (slit width 20 eV for higher magnifications; Gatan, Inc.), and a 4 k × 4 k K2 Summit direct detector (Gatan, Inc.) in counting mode.

First, cellular areas containing the fluorescent bodies of interest in suitably-thin areas that were typically 200-500 nm thick were located in the TEM using methods described previously. Tilt series were then recorded of these areas using UCSF Tomography (*43*) or SerialEM (*44*) software at a magnification of 27,500× (Polara) and 34,000× (Polara). This corresponds to a pixel size of 3.712 Å (MEFs) or 3.260 Å (U2O2), respectively, at the specimen level and was found to be sufficient for this study. Each tilt series was collected from −60° to +60° with an increment of 1° in an automated fashion at 4–10 μm underfocus. The cumulative dose of one tilt-series was between 80 and 150 e−/Å^2^. The tilt series was aligned and binned by 4 into 1k × 1k using the IMOD software package (*45*), and 3D reconstructions were calculated using the simultaneous reconstruction technique (SIRT) implemented in the TOMO3D software package (*46*), or weighted back projection using IMOD. Noise reduction was performed using the non-linear anisotropic diffusion (NAD) method in IMOD (*45*), typically using a K value of 0.03–0.04 with 10 iterations.

### Segmentation and isosurface generation

Segmentation and isosurface rendering were performed in Amira (Thermo Scientific). For lower resolution tomograms from MEFs cells, the segmentation was done all manually using the brush, lasso, and thresholding tools combined with interpolation and surface smoothing. For higher resolution tomograms from U2-OS cells, the ER, IRE1α subdomains and vesicles were segmented manually. Cytoskeletal components and ribosomes were segmented using the TomoSeg CNN module in EMAN2 (*33*).

### Conventional electron microscopy

HEK293 stable cells expressing IRE1-GFP, unstressed or treated with tunicamycin, were fixed with 2.5 % glutaraldehyde, 2% paraformaldehyde in 0.1 M Sorenson’s phosphate buffer (PB), pH 7.4, for 2 h at room temperature. After storage in 1% PFA, 0.1 M PB at 4 °C for about 1 week and rinsing in 0.1 M PB, the cells were postfixed with 1 % OsO_4_ and 1.5 % K_3_Fe(CN)_6_ in 0.07 M PB, stained *en bloc* in aqueous 0.5 % uranyl acetate, dehydrated in acetone and embedded in Epon. Ultrathin plastic sections were stained with uranyl acetate and lead citrate and examined in a JEOL JEM-1010 electron microscopy. Quantification of the diameters of IRE1a subdomain tubes and ER cisterns was performed using Fiji software (*47*).

### Immuno-electron microscopy

HEK293 cells stably expressing IRE1-GFP, untreated or treated with tunicamycin, were fixed using 4 % PFA in 0.1 M PB for 2 h at room temperature, then overnight at 4 °C. Subsequently, the fixation was continued by replacing the initial fixative with 1 % PFA in 0.1 M PB for several days. The samples were then rinsed in PBS, blocked with 0.15 % glycin in PBS, scraped in 1% gelatin in PBS, pelleted, and embedded in 12 % gelatin. Small blocks of pellet were cryoprotected with 2.3 M sucrose, mounted on aluminum pins and frozen in liquid nitrogen. Ultrathin cryosections were cut at −120 °C, placed on copper carrier grids, thawed and immunolabeled as previously described (*48*). In brief, the sections were incubated with blocking buffer containing fish skin gelatin (Sigma-Aldrich, G7765) and acetylated BSA (Aurion, 900.022) and immunolabeled with biotinylated goat anti-GFP antibody (Rockland, 600-106-215) at 1:300, followed by rabbit anti-biotin antibody (Rockland, 100-4198) at 1:10000. Subsequently, the sections were incubated with Protein A-conjugated 10 nm gold particles (Cell Microscopy Core, University Medical Center Utrecht, the Netherlands), stained with uranyl acetate followed by a methylcellulose-uranyl acetate mixture, and examined in a JEOL JEM-1010 electron microscope. Quantification of the diameters of IRE1a subdomain tubes and ER cisterns was performed using Fiji software *(47)).*

### Subtomogram averaging

In U2OS cells each tilt series was collected from −60° to +60° with an increment of 1°in an automated fashion using SerialEM at 4–6 μm underfocus. The image pixel size used for subtomogram averaging was 6.52 Å (binned by 2). Subsequent subtomogram averaging was performed by the EMAN2 tomography pipeline *(49).* Initially unbinned tilt-series were automatically aligned and reconstructed using EMAN2. In total, 3 cryo-tomograms were generated to provide a sufficient number of particles for further processing. Particles were picked using EMAN2 particle picking software using a box size of 56×56×56 pixels. Briefly model points were placed every 10 nm along the length (approximately one helical turn), of the oligomer present inside the membrane tube. Particles of various orientations were picked, including top-views and side-views. In total 653 model points were picked. CTF estimation and correction was performed by EMAN2. An initial model was then generated in C1 with all 653 particles in 5 iterations. The iteration 5 map was aligned to the symmetry axis and was used as an initial model for subtomogram refinement using C2 and D2 symmetry. The first iteration D2 map generated in subtomogram refinement was used as a reference for a sub-tilt refinement step using helical symmetry in 5 iterations. The helical symmetry parameters were as follows; C symmetry = 2, rotation about the Z axis = 45°, nsym = 2.5, and tz = 5. The map produced by iteration 5 includes 80% of the best aligned particles and is shown in figure S10. To focus on one strand of the helix, an automatic mask was generated to improve alignment in 5 iterations. The one-stranded map produced by iteration 5 includes 80% of the best aligned particle and is shown in figure 4. The particles were split into two subsets and resolution is measured by the Fourier shell correlation of these two density maps. The correct hand of the final map was determined by EMAN2. The particles were mapped back into the cryo-tomogram by EMAN2 using a pKeep of 0.6.

### Model prediction

A schematic illustrating the model building process has been included in Fig. S11. SWISS-MODEL was used to predict the human active form of IRE1α, based on the human sequence and the *S. cerevisiae* IRE1-LD crystal structure (PDB:2BE1) as a template. To prepare the template for SWISS-MODEL, the missing loops in the crystal structure were built using Modeller in Chimera (*50*).

### Dimer-dimer interface generation

Two predicted model IRE1α dimers were superimposed onto the *S. cerevisiae* IRE1-LD crystal structure dimer-dimer in Chimera. The *S. cerevisiae* IRE1-LD crystal structure dimer-dimer was then omitted, leaving a gap between the two predicated human IRE1α dimers. A new interface was modelled in Chimera by translating one dimer along one axis until the gap was filled. To accommodate the new dimer-dimer interface in the double-helical map, the angle between the two dimers was made more acute to approximately 45° (the rise of the helix). The resulting dimer was then placed into the map using the fit-to-map function in Chimera. The remainder of the helix was then built whilst maintaining the same dimer-dimer interface throughout the helix.

### Quantifications

IRE1α subdomain tube diameters were measured by drawing lines between the center of each membrane density perpendicular to tube membranes. Regions less than 30 nm away from a junction are excluded. Tube diameters for 12 representative tomograms were quantified as 409 measurements. Distributions of immunogold particles shown in Fig. 2 were generated by manually extracting the X-Y coordinates of each gold particle and using a python script to measure all pairwise distances and plotting the distance distribution. Line intensity plots shown in Fig. 3 were generated by averaging and aligning multiple line plots across each subdomain cross section. The resulting averaged plates were subsequently aligned and averaged as shown in Fig. 3C. Quantification of radii of curvature was performed by dividing IRE1α subdomain tubes clearly visible in XY planes into tiling 25 nm segments, excluding junctions. Circles with diameters in increments of 50 nm were then manually fitted to each segment to yield estimates of the radius of curvature. N = 274.

### Calculations

Calculations of IRE1 domain concentration assumes IRE1α subdomain tubes to be perfect cylinders of 28.3 nm diameter measured from membrane centers with a membrane thickness of 37.5 Å (*51*). Each 100 nm segment of such cylinder contains 139 IRE1-LD monomers, approximated from a pitch of 173 Å with 24 monomers per turn. The volume occupied by IRE1α-LD is calculated by using a glycosylated MW of 49196 g/mol and a density of 1.35 g/mL (*52*), yielding a volume of 8.41 × 10^3^ nm^3^ occupied by 139 IRE1-LD monomers and 3.87 × 10^4^ nm^3^ void volume. In this void volume, a single substrate molecule has a concentration of 43 μM. IRE1 density fold change upon stress are approximated from 139 monomer per 0.008890 μM^2^ and contrasting to earlier calculations (*25*). IRE1-LD concentration is extracted from the molarity of 139 units as 2.31 × 10^−22^ mole / 4.7144* 10^−20^ L. IRE1 cytosolic domain concentrations are estimated from the approximate volume experienced by this domain (1.05×10^6^ nm^3^ which assumes the domain extends 44 nm from the membrane due to a stretched cytosolic linker + KR domain height from crystal structures). The fold compaction for the KR domain from full distribution along cylindrical tubes to a flat ER membrane compares the experienced volume between a hollow cylinder as above and a column trapezoid with a volume of 2.49×10^5^ nm^3^, which yielded the ~4 fold increase.

---

**Fig. S1.**
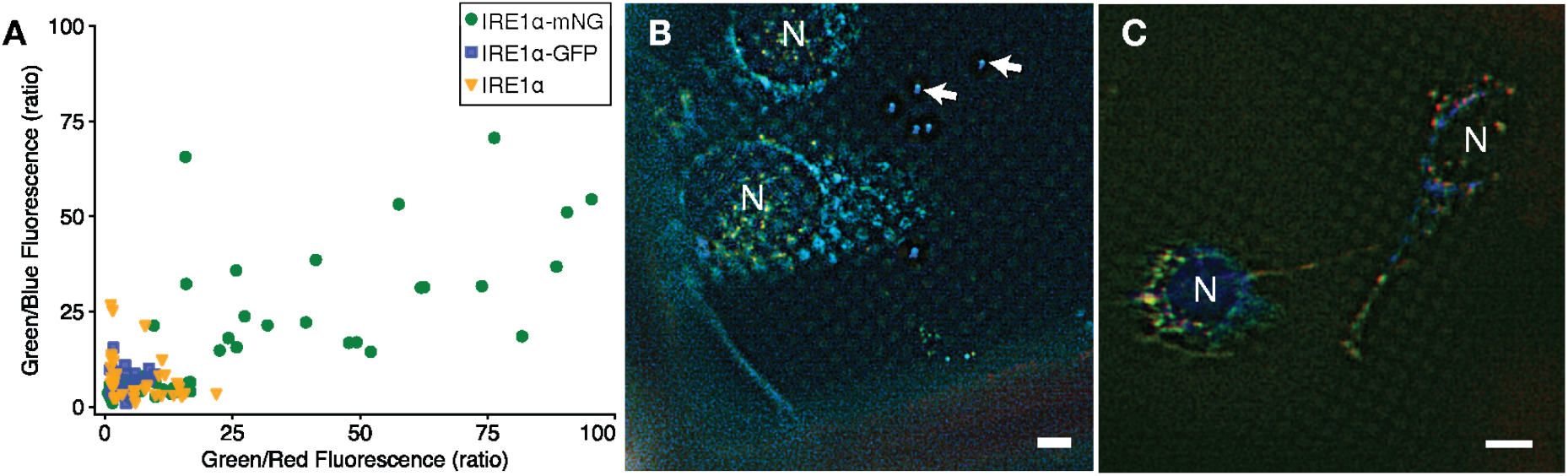
Fluorescent profile across cell lines expressing different IRE1α constructs. (A) Scatter plot of fluorescent profiles for bright spots observed at liquid nitrogen temperature for stressed cells expressing IRE1α-mNG, IRE1α-GFP or WT IRE1α. The ratios of Green/Red and Green/Blue fluorescence intensity taken at the same exposures across channels are used to identify specific IRE1α signal. Representative cryo-fluorescent light microscope images of stressed MEFs-IRE1α-GFP (B) and MEFs-IRE1α (C; lacking fluorescent protein) cells grown directly on coated Quantifoil grids imaged at 1 second exposure in BFP, GFP and mCherry channels. Cells were treated with Tm for 2 hours prior to cryo-preservation and imaged in liquid nitrogen. Nuclei are indicated with “N.” White arrows: 500 nm fluorescent nanosphere. Scale bars are 6 μm.

**Fig. S2.**
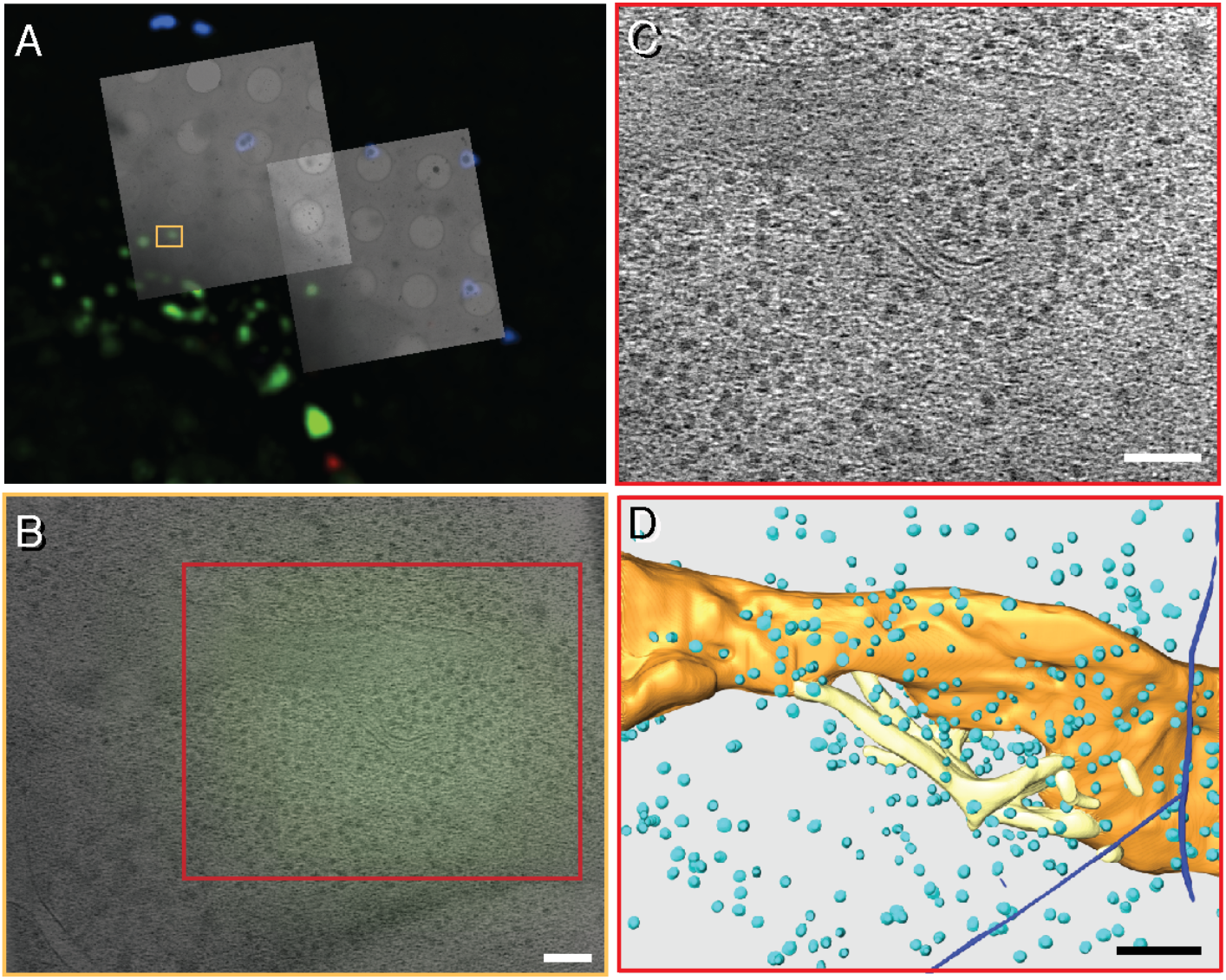
Target identification by correlation and tomographic reconstruction in intact MEFs cells expressing IRE1α-mNeonGreen. (A) Overlay of multi-channel cryo-LM images with low magnification (3000X) cryo electron micrographs of regions containing IRE1-mNeonGreen foci formed after 2 hours ER stress induction. Blue fluorescent spheres visible by both light and electron microscopy were used to align the images in × and Y directions prior to targeting cell regions for high magnification tilt series. (B) Correlation of reconstructed tomogram with fluorescent signal reveals sub-regions with IRE1 foci in an additional example from the same cell depicted in Fig. 1B-F. (C) A z slice showing thin membrane tubes within regions colocalizing with IRE1-mNeonGreen signal. (D) manual 3D segmentation for tomogram in C. Orange: normal ER membrane. Yellow: thin ER membranes at IRE1-mNeonGreen regions. Teal: ribosomes. Blue: cytoskeletal elements. Scale bars: 100 nm.

**Fig. S3.**
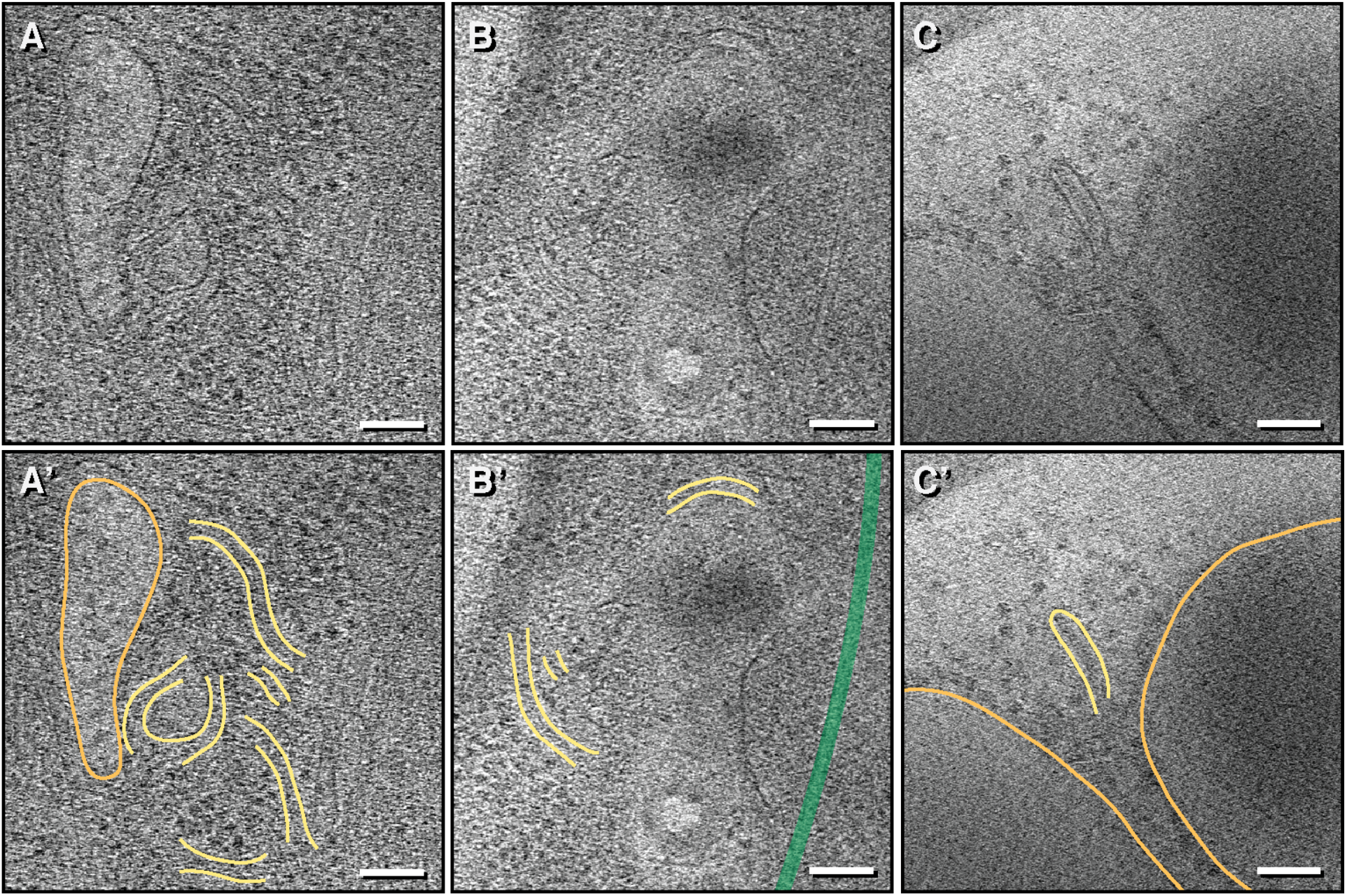
Additional example of IRE1α-mNG fluorescence correlated to narrow membrane tubes in stressed MEFs-IRE1α-mNG cells. (A-C) Representative z slice of corresponding sub-regions correlated with IRE1α-mNG signal in 3 different examples obtained in stressed cells expressing IRE1α-mNG. Scale bar = 100 nm. (A’-C’) Segmentation of membranes and structures observed in tomograms where yellow = thin ER tubes of IRE1α subdomains, orange = membrane of ER sheet/tubules, and green = cytoskeletal components.

**Fig. S4.**
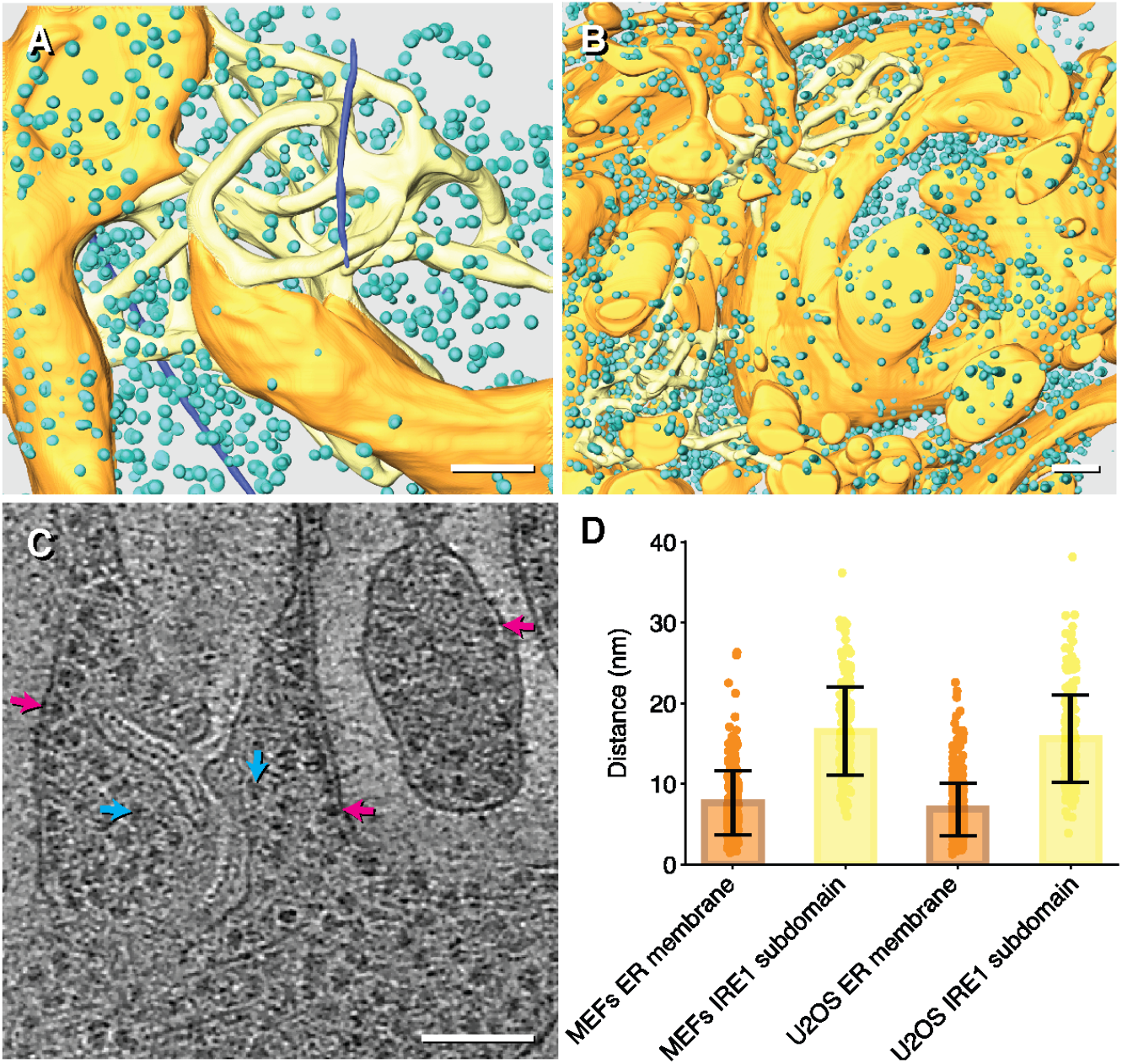
IRE1α subdomain tubes are devoid of bound ribosomes. (A) Segmentation of MEFs tomogram corresponding to Fig. 1D but showing ribosomes and cytoskeletal segmentation. Scale bar = 100 nm (B) Segmentation of U2OS tomogram with larger cell region corresponding to Fig 1F with ribosomes and cytoskeletal components shown. Scale bar = 100 nm. (C) An example z slice of IRE1α subdomain tube and adjacent ER sheet/tubules. Cyan arrows point to ribosomes near IRE1α subdomains; Magenta arrows indicate ribosomes bound to ER membranes. Scale bar = 100 nm. (D) Plot of distance between ribosomes and membranes comparing ER sheet/tubules membranes and IRE1α subdomain membranes for tomograms in A and B. Error bar is standard deviation. N = 218, 180, 215, 519.

**Fig. S5.**
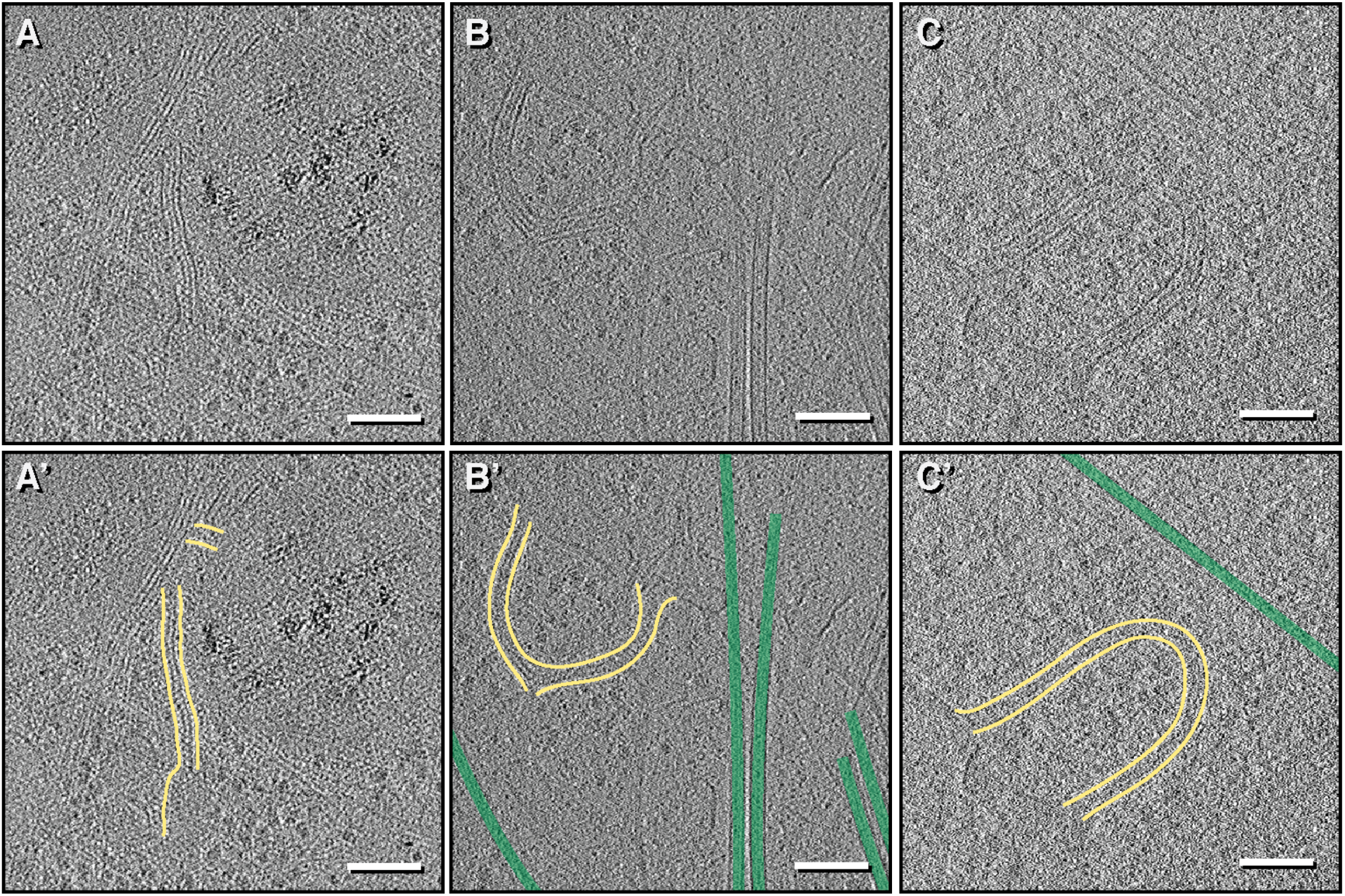
Additional example of IRE1α fluorescence correlated to narrow membrane tubes in stressed U2OS cells expressing fluorescently tagged IRE1α. (A-B) Representative z slice from two additional examples obtained in stressed U2OS-IRE1α-mNG cells expressing fluorescently tagged IRE1α. (C) representative Z slice from an example obtained from U2OS-IRE1α-mRuby cell line where IRE1α is tagged at same location with an mRuby3 red fluorescent protein instead of mNeonGreen. Observation of narrow membrane tubes with specific red fluorescence using mRuby3 supports that bright fluorescent foci are indeed IRE1α foci rather than autofluorescent signal. (A’-C’) Segmentation of membranes and structures observed in tomograms where yellow = thin ER tubes of IRE1α subdomains and green = microtubules. Scale bar = 100 nm.

**Fig. S6.**
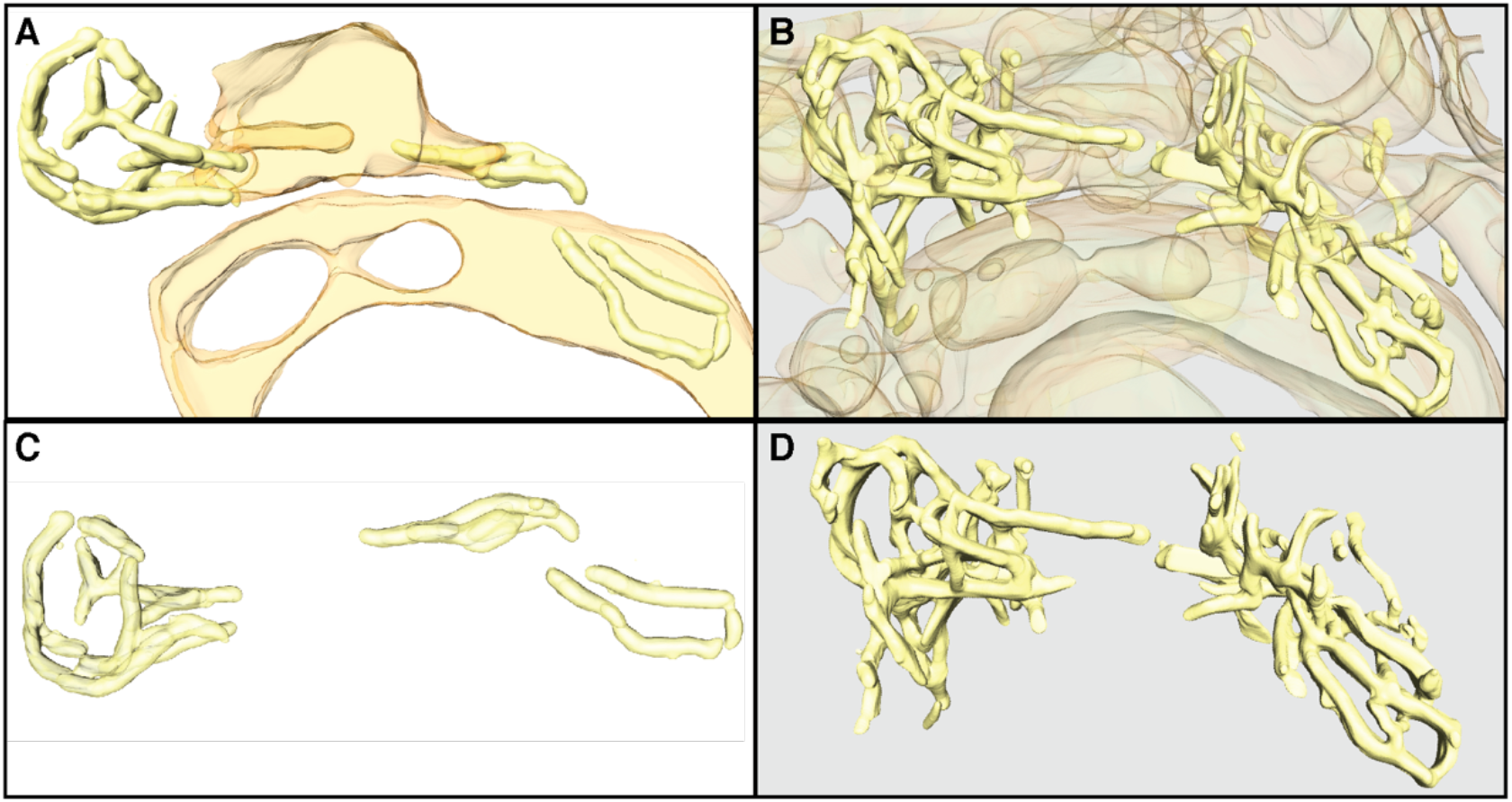
Comparison of machine-learning and manual segmentation of U2OS tomogram. (A) and (C) are isosurfaces generated from EMAN2 CNN segmentation where general ER structures are shown in (A) and only the narrow RE1α subdomain tubes are shown in C. (B) and (D) are comparable views with smoothed isosurfaces generated manually using manual segmentation tools in Amira. A lot more structures are recognized by human eyes than are annotated by the neural network, especially at junctional regions. The manual segmentation in (B) and (D) therefore has higher degree of connectivity compared to (A) and (C).

**Fig. S7.**
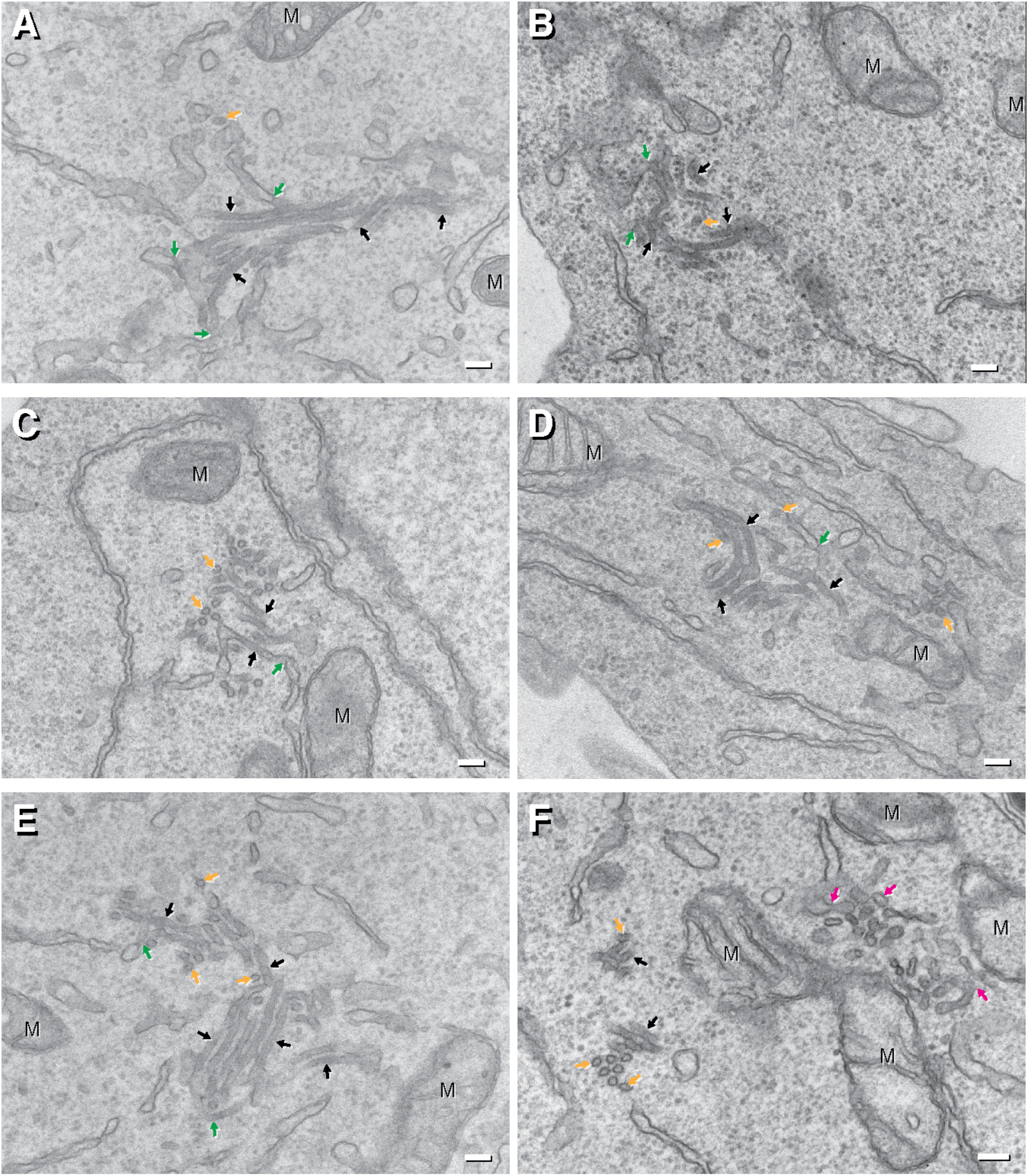
Rare instances of IRE1α subdomain can be observed in stressed cells expressing inducible IRE1α-GFP by conventional EM. (A-F) Micrographs of thin sections of Epon-embedded cells. Thin membrane structures with diameters of 30 ± 3 nm (error: standard deviation. N=89) can be observed as rectangular longitudinal (black arrows) and circular end-on cross-sections (orange arrows). These narrow membranes are connected to larger ER structures (diameters of 53 ± 20 nm, N=90) at junctions indicated by green arrows. Intriguingly, the lumens of these structures appear darker than surrounding ER lumen, indicative of higher protein density. Scale bar = 100 nm. M = mitochondria. These extremely narrow membrane tubes are morphologically distinct from ER exit sites (magenta arrows in F), which are finger-like protrusions with diameters of 35 ± 7 nm (N=20) consistent with ER-derived vesicles.

**Fig. S8.**
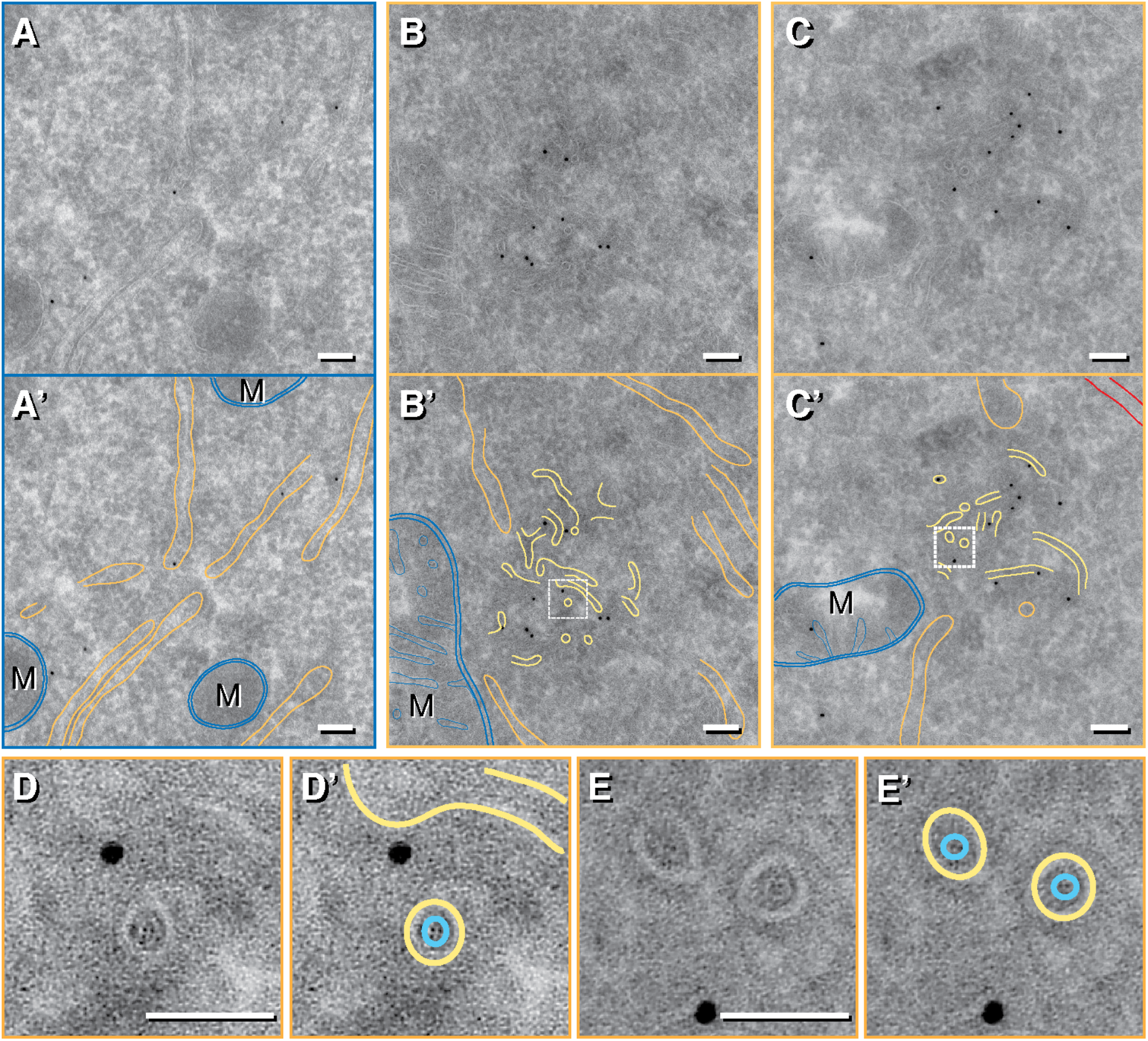
Additional example of visualization of IRE1α subdomain by immunogold EM. (A, A’) Micrograph of cell expressing IRE1α-GFP epitope but not stressed, showing disperse and sparse immunogold particles labeling general ER tubules/sheets segmented in orange. M = mitochondria are segmented in blue. Scale bars = 100 nm. (B, B’, C, C’) Micrograph of immunogold-labeled stressed cells expressing IRE1α-GFP epitope. Large clusters of gold particles localize to regions with narrow membrane structures (segmented in yellow). These narrow tubes have an averaged diameter of 26 ± 2 nm (error: standard deviation; N = 108) compared to surrounding ER structures segmented in orange with an averaged diameter of 51 ± 18 nm (error: standard deviation; N = 133). M = mitochondria are segmented in blue. Plasma membranes seen in C are segmented in red. Scale bars = 100 nm. Regions within white squares in B’ and C’ are enlarged in D and E, respectively. End-on cross sections of thin IRE1α subdomain tubes appear as two roughly concentric rings, consistent with helical protein density within the lumen (cyan density). Scale bar = 50 nm.

**Fig. S9.**
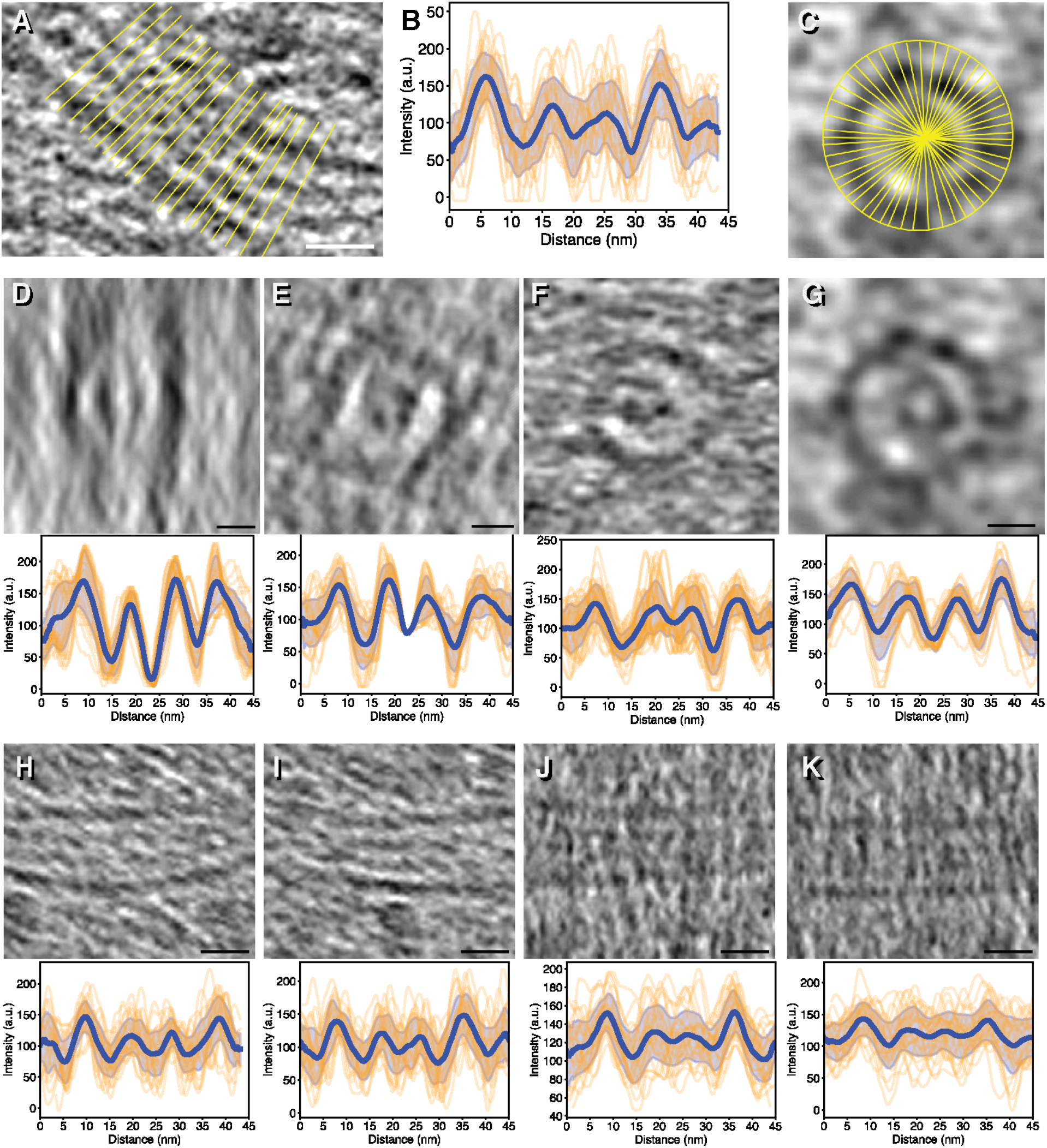
IRE1α subdomain tubes in MEFs cells contain lumenal protein density. (A) Example of how line profiles were drawn for IRE1α subdomain longitudinal cross sections and resulting averaged line plot (B) showing four distinct peaks. (C) Example of how line profiles were drawn for IRE1α subdomain end-on cross section depicted in Fig. 3B. (D-K) IRE1α subdomain cross sections and resulting averaged line plots used to generate plot of averaged profiles and measure diameters shown in Fig. 3C. Scale bars = 20 nm.

**Fig. S10.**
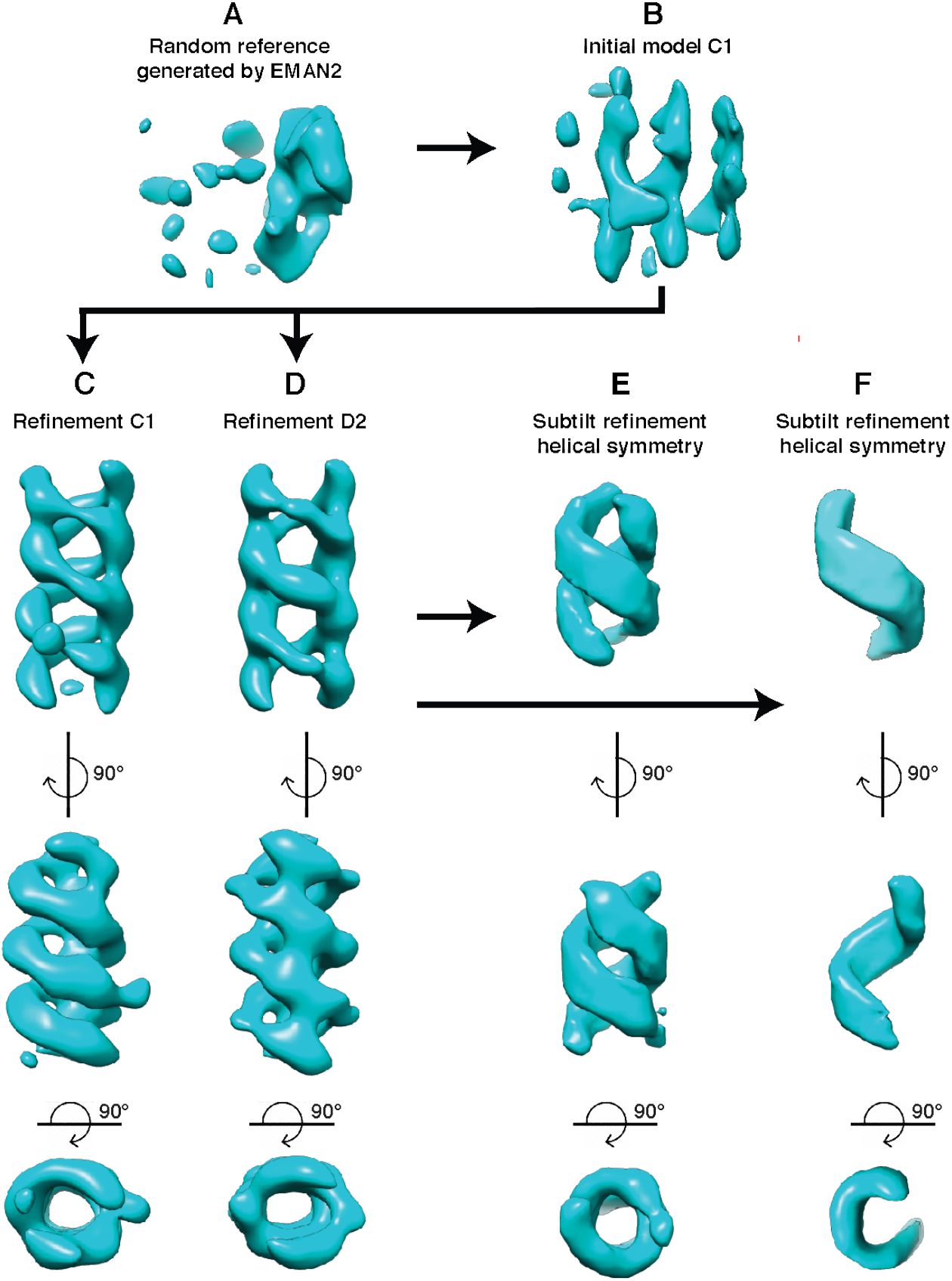
Subtomogram average workflow. (A) EMAN2 generated random reference. (B) Initial C1 model. (C) Refinement of the initial model in C1. (D) Refinement of the initial model in D2. This D2 average was used as a reference for all the following subtilt refinements, (E) helical symmetry (F) helical symmetry focused on one strand of the double-helical filament.

**Fig. S11.**
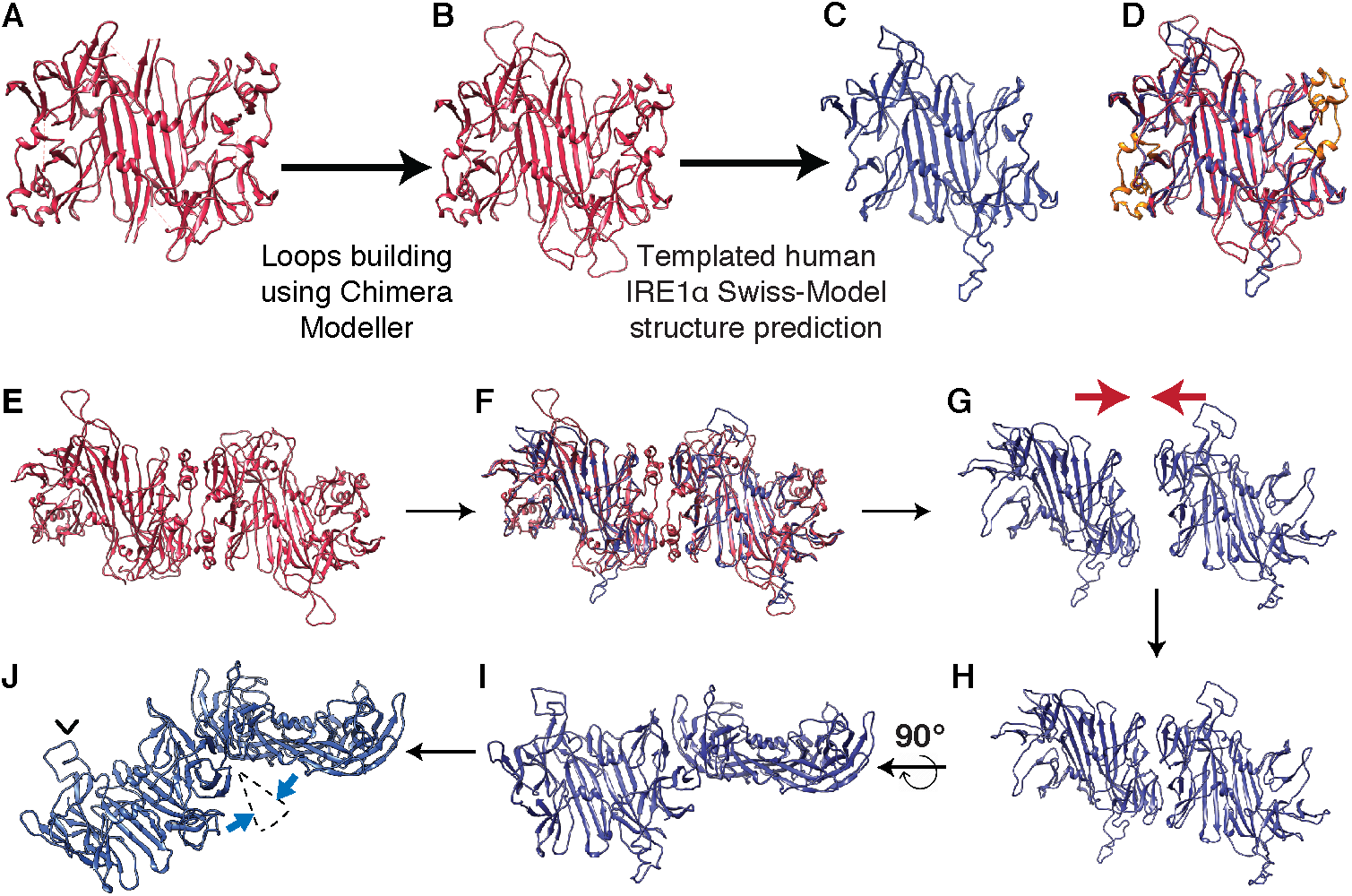
Generation of a modeled human active IRE1α-LD and dimer-dimer interface. (A-D) SWISS-MODEL homology modeling of the active human IRE1α LD using the *S. cerevisiae* IRE1 lumenal crystal structure (PDB ID: 2BE1) as a template. (A) 2BE1. (B) 2BE1 with loops built using Modeller in Chimera. (C) Human IRE1α lumenal structure predicted using the template generated in B. (D) Superimposing of 2BE1 (B; with loops) and modeled active human IRE1α lumenal domain (C) reveals a helical bundle present in B but not in C, shown in orange. (E) The *S. cerevisiae* IRE1 lumenal crystal structure (2BE1) dimer. (F) Two modeled active human IRE1α-LDs were superimposed onto the dimer in E. (G) Same superimposed human IRE1α-LDs as F, with the *S. cerevisiae* dimer removed. Red arrows highlight translational shift imposed on the two monomers to eliminate gap, resulting in a new interface shown in H. (I) Same as H, rotated 90° around the axis of the interface hinge. (J) Hinge angle decreased as indicated by blue arrows to accommodate the curvature of the sub-tomogram average maps.

**Fig. S12.**
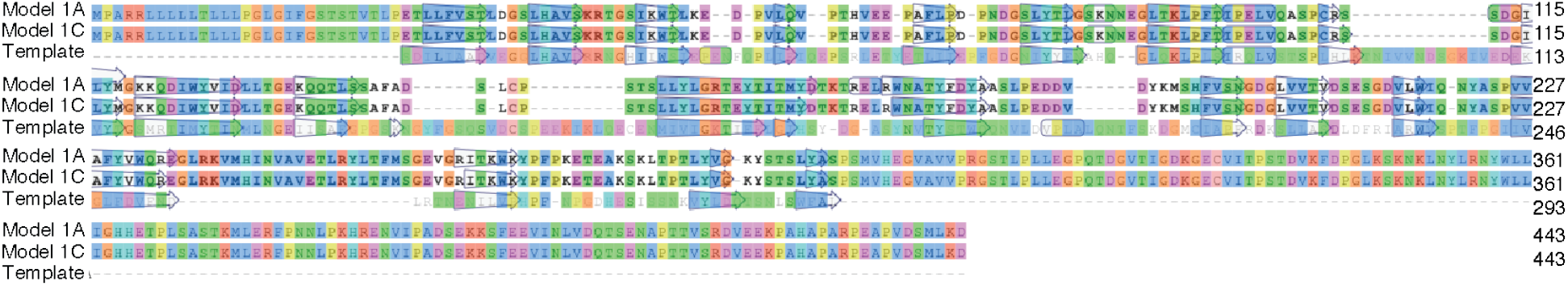
Model-Template sequence and secondary structure alignment in DSSP format for the SWISS-MODEL homology modeling of the active human IRE1α LD shown in S11.

**Fig. S13.**
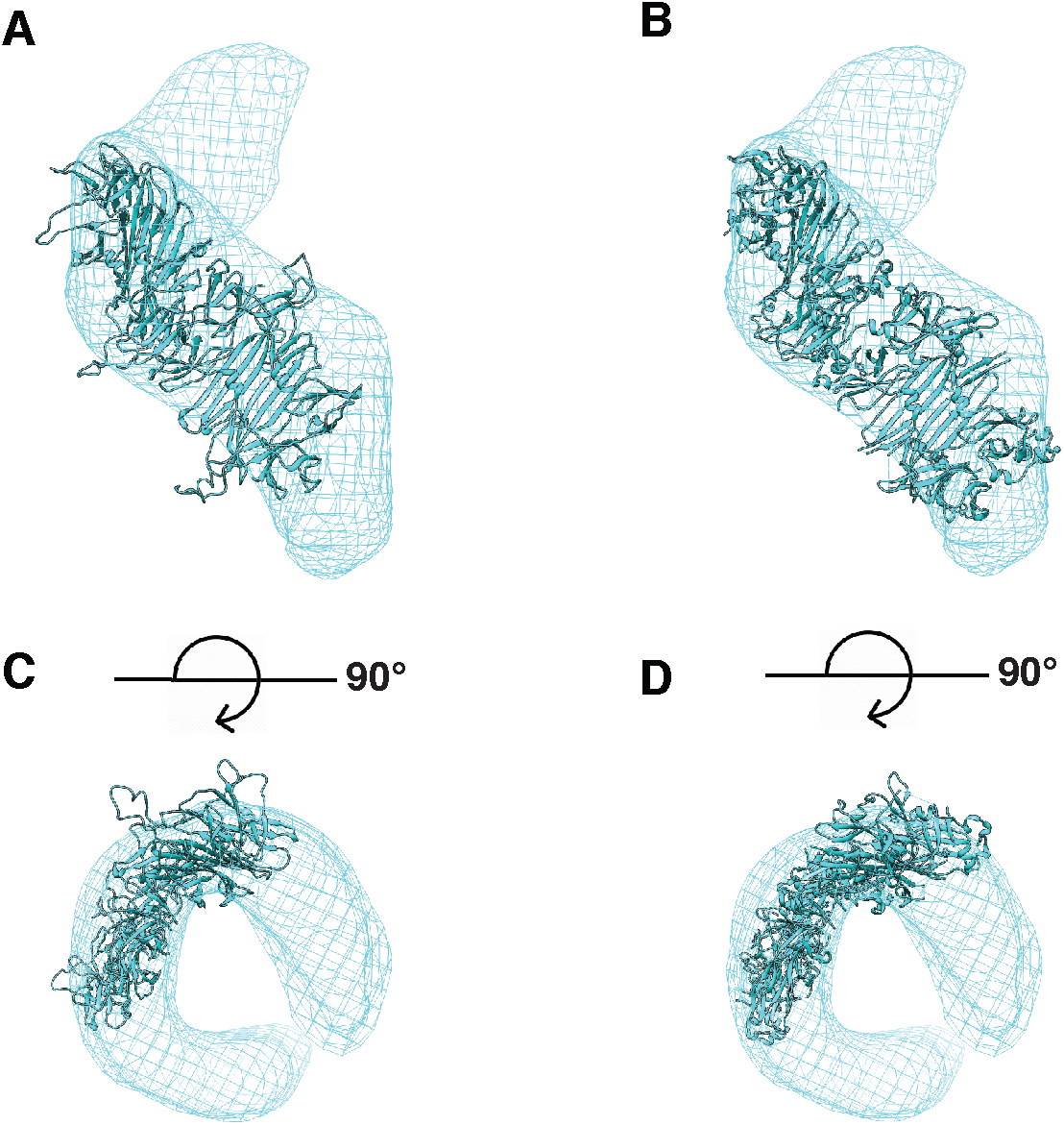
Steps used to build double-helical model. (A) The dimer-dimer in the human active IRE1α-LD fitted into map as a rigid body using fit in map function in Chimera. (B) Generation of an altered dimer-dimer interface based on the rise and twist of the helix was fitted into map as a rigid body using fit in map function in Chimera. (C, D) Same views as A and B, respectively, rotated around the x-axis by 90°.

**Fig. S14.**
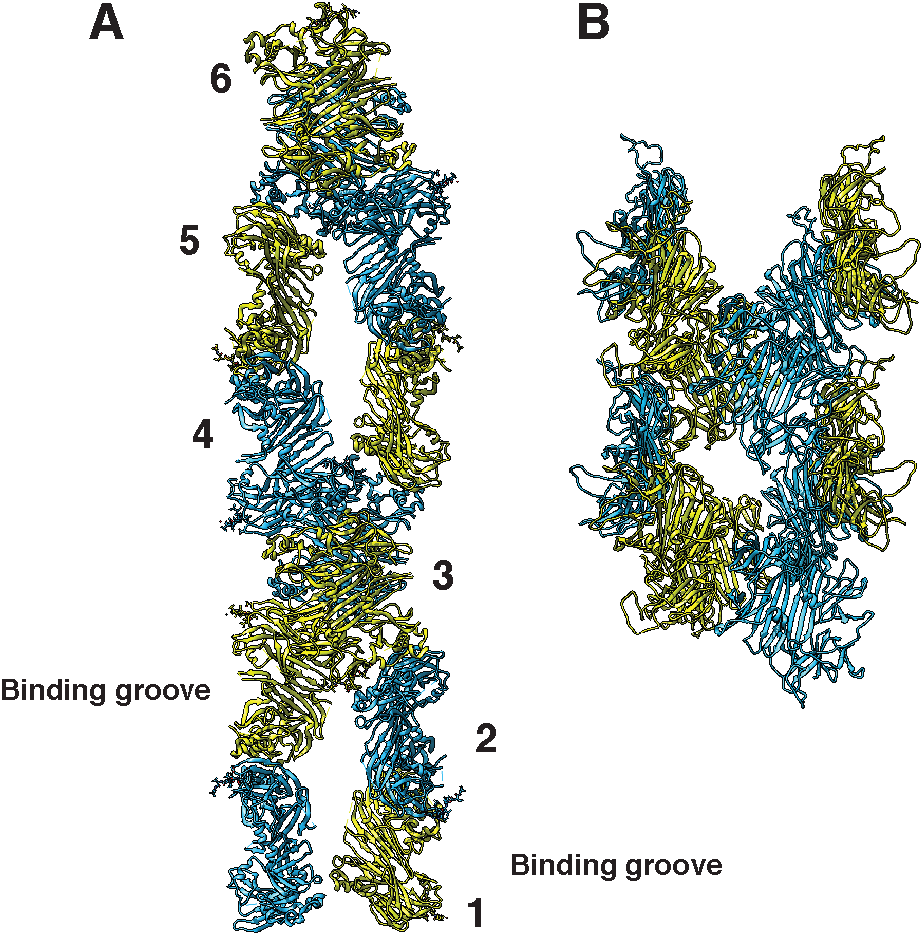
Comparison of *S. cerevisiae* IRE1-LD 2BE1 and modeled active human IRE1α-LD oligomers. (A) Double helical oligomer found in the crystal structure of the *S. cerevisiae* IRE1-LD with 12 dimers per double-helical turn and the predicted unfolded protein binding groove facing outward. Numbers indicate individual dimers running along one helical filament. (B) the human IRE1α-LD double helix modelled and fitted to the sub-tomogram averaging map.

**Fig. S15.**
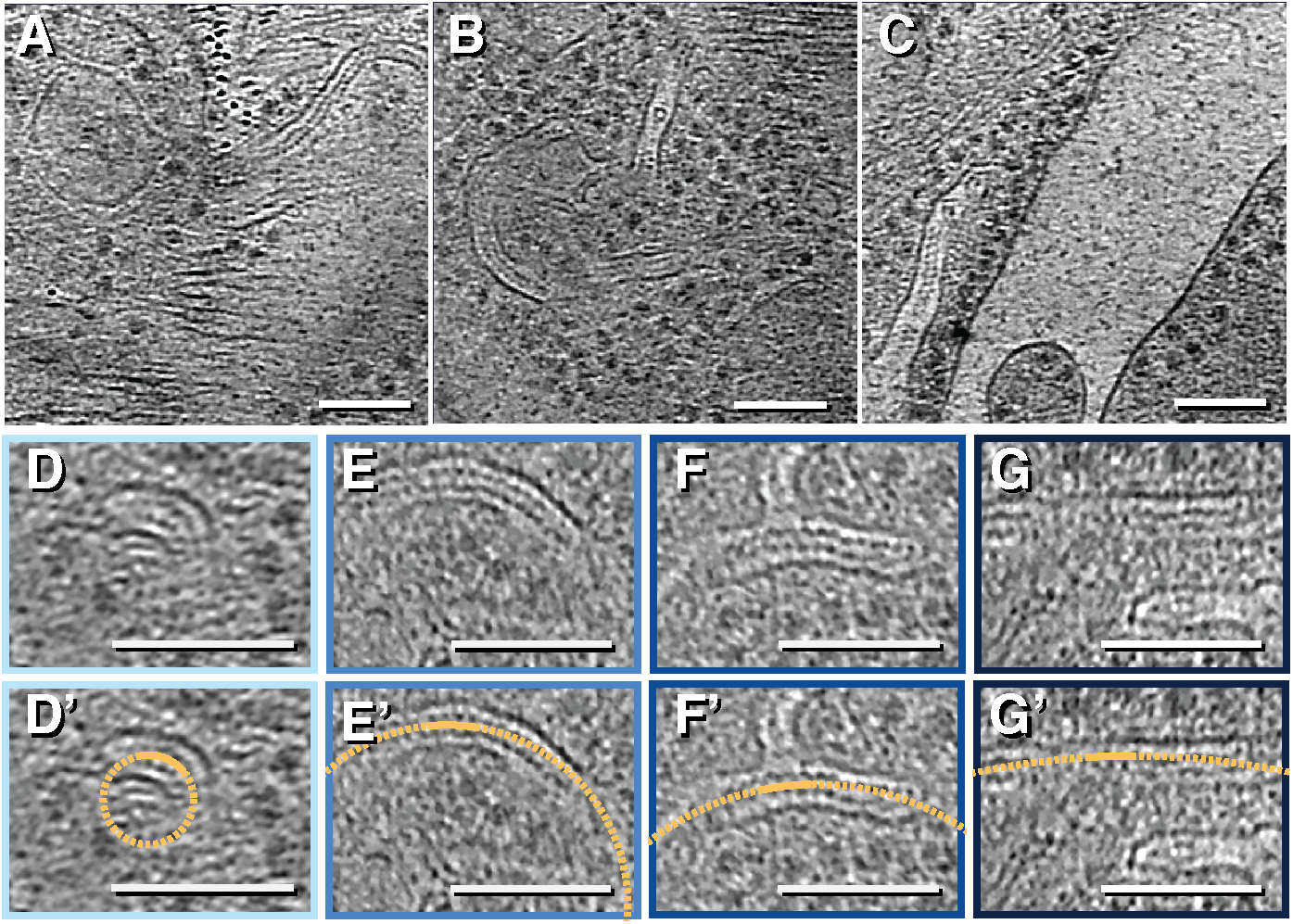
Example range of curvatures observed for IRE1α subdomains. (A-C) Example regions from U2OS-IRE1-mNG cells showing IRE1α subdomain tubes with varying degrees of curvature. Cropped regions exhibiting radii of curvature (ROC) ranging from 25-75 nm (D), 75-125 nm (E), 125-175 nm (F), and >175 nm (G) quantified and plotted in Fig. 4D. The outline color matches the bars in Fig. 4D, organized as increasing darkness corresponding to increasing ROC. A sample local curvature fit circle (dash orange circle) with ROC of 25 nm (D’), 100 nm (E’), 150 nm (F’) and 400 nm (G’) are shown for representative 25nm segments (solid orange arcs) within the view shown. All scale bars = 100 nm.

**Fig. S16.**
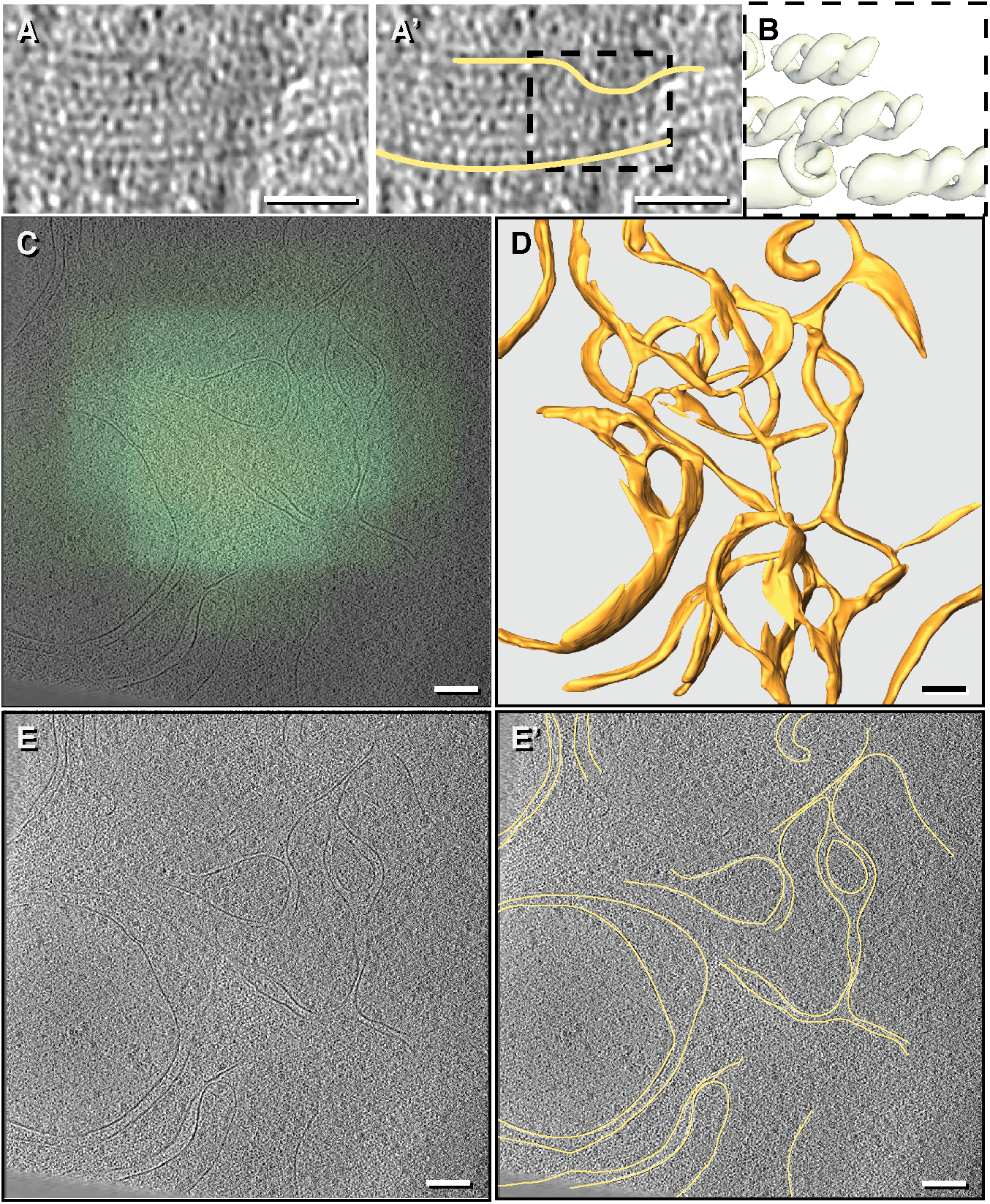
Anomaly in IRE1α subdomains structures. (A, A’) An example of an IRE1α subdomain tube containing multiple juxtaposed IREα-LD filaments not separated by membrane (yellow). Scale bar = 20 nm (B) An isosurface of the averaged density mapped back into this region. (C) Overlay of cryo-light microscopy image and a representative slice of the reconstructed tomogram showing colocalization of IRE1α-mNG foci to cell sub-regions with complex membrane topology and irregular inter-membrane diameters. Scale bar = 100 nm. (D) Segmentation of this putative precursor of IRE1α subdomains with regular tube diameter, showing thin tubes connected by 3-ways junctions and enrichment of ER-fenestrations. (E, E’) An example z slice of the membrane structures observed. Scale bars = 100 nm.

**Movie S1.**

IRE1α-mNeonGreen foci in stressed Mouse Embryonic Fibroblasts localize to narrow IRE1α subdomain tubes.

**Movie S2.**

IRE1α-mNeonGreen foci in stressed U-2 OS cells localize to narrow IRE1α subdomain tubes. ER tubules and sheet membranes are segmented in orange. IRE1α subdomain tubes are shown in yellow. Other cellular membranes are shown in blue. Within the narrow IRE1α subdomain tubes, lumenal protein densities with helical feature can be observed by stepping through the 3D tomogram. The lumenal double helices, resolved by subtomogram averaging, are flexible and are consistent with modeled helical arrays of human IRE1α lumenal oligomers.

**Movie S3.**

An additional example of an IRE1α-mNeonGreen focus in U-2 OS cells.

